# Phosphorylation-Dependent Association of WRN with RPA is Required for Recovery of Replication Forks Stalled at Secondary DNA Structures

**DOI:** 10.1101/2023.08.08.552428

**Authors:** Alessandro Noto, Pasquale Valenzisi, Federica Fratini, Tomasz Kulikowicz, Joshua A. Sommers, Flavia Di Feo, Valentina Palermo, Maurizio Semproni, Marco Crescenzi, Robert M. Brosh, Annapaola Franchitto, Pietro Pichierri

## Abstract

The WRN protein mutated in the hereditary premature aging disorder Werner syndrome plays a vital role in handling, processing, and restoring perturbed replication forks. One of its most abundant partners, Replication Protein A (RPA), has been shown to robustly enhance WRN helicase activity in specific cases when tested *in vitro*. However, the significance of RPA-binding to WRN at replication forks in vivo has remained largely unexplored. In this study, we have identified several conserved phosphorylation sites in the acidic domain of WRN that are targeted by Casein Kinase 2 (CK2). Surprisingly, these phosphorylation sites are essential for the interaction between WRN and RPA, both *in vitro* and in human cells. By characterizing a CK2-unphosphorylatable WRN mutant that lacks the ability to bind RPA, we have determined that the WRN-RPA complex plays a critical role in fork recovery after replication stress whereas the WRN-RPA interaction is not necessary for the processing of replication forks or preventing DNA damage when forks stall or collapse. When WRN fails to bind RPA, fork recovery is impaired, leading to the accumulation of single-stranded DNA gaps in the parental strands, which are further enlarged by the structure-specific nuclease MRE11. Notably, RPA-binding by WRN and its helicase activity are crucial for countering the persistence of G4 structures after fork stalling. Therefore, our findings reveal for the first time a novel role for the WRN-RPA interaction to facilitate fork restart, thereby minimizing G4 accumulation at single-stranded DNA gaps and suppressing accumulation of unreplicated regions that may lead to MUS81-dependent double-strand breaks requiring efficient repair by RAD51 to prevent excessive DNA damage.

## INTRODUCTION

The Werner’s syndrome protein (WRN) is one of the five conserved RECQ helicases in humans and is the protein mutated in the rare genetic disease Werner’s syndrome (WS) (*1–4*). WRN plays critical roles in the maintenance of genome integrity, in particular during DNA replication as evidenced by several characteristic phenotypes of WS patient-derived or WRN-depleted cells (*1*, *5*, *6*). During DNA replication, WRN is required for multiple functions including DSBs avoidance, proper fork recovery and replication of fragile sites, end-processing of reversed and of collapsed forks (*1*, *6*). More recently, WRN was found to participate to fork protection in BRCA2-deficient cells, limit R-loop-associated DNA damage, and assist in the stabilisation of microsatellites (*7–9*).

The genome caretaker functions carried out by WRN during DNA replication involves multiple protein-protein interactions with other crucial factors implicated in DNA replication under stressed conditions (*5*, *6*). One of the most abundant WRN interactors playing key roles in response to perturbed replication is RPA (*10*). RPA heterotrimer is the major human single-strand DNA (ssDNA) binding protein, which recognises ssDNA formed during DNA replication or repair also acting as a scaffold for other factors implicated in the response to perturbed replication (*11–13*). WRN protein binds to the N-terminal domain of RPA1 through its acidic domain *in vitro* and colocalises with RPA at replication foci in human cells (*10*, *14–17*). Although, *in vitro*, the WRN-RPA association stimulates WRN helicase activity on branched substrates mimicking stalled or reversed replication forks (*10*, *14*, *17–20*), it is unknown which of WRN’s different functions requires the interaction with RPA in response to perturbed replication.

Here, we identified multiple phosphorylation sites in the WRN acidic domain that are targeted by CK2 and essential to drive the interaction of WRN with RPA *in vitro* and in cells. We used the WRN 6A mutant, which is unphosphorylable by CK2 and defective in RPA-binding, as a tool to assess the functional relevance of RPA-binding during the response to perturbed replication. This mutant contains Ser/Thr to Ala substitutions at all six sites targeted by CK2, but retains the ability to relocalise to ssDNA like the wild-type WRN. Characterisation of the response to DNA replication perturbation of cells expressing the WRN 6A mutant as compared with cells expressing the wild-type WRN revealed that RPA-binding is not involved in the WRN-dependent end-processing occurring at stalled or collapsed forks or in limiting formation of DSBs. By contrast, WRN-RPA interaction is required to properly restart stalled forks limiting accumulation of parental ssDNA gaps and allowing efficient clearance of G4s DNA. When the WRN-RPA interaction is negatively affected or the WRN helicase is inhibited, MUS81 contributes to remove G4s producing DSBs downstream the formation of MRE11-dependent gaps. RAD51-dependent repair is subsequently required to limit accumulation of DNA damage. Together these findings help to clarify the function of WRN binding to RPA for a proper response to replication stress at G4s.

## RESULTS

### The acidic domain of WRN is phosphorylated by CK2

The high-affinity RPA-binding site of WRN is located in its acidic domain (*17*) and a cluster of high-ranking putative CK2 phosphorylation sites can be identified in this domain (Figure 1a). To assess if the acidic domain of WRN is targeted by CK2, we first expressed in bacteria the GST-tagged WRN N-terminal fragment or, as a control, the GST alone and phosphorylated the corresponding GST fusion protein *in vitro* using recombinant CK2 (Supplementary Figure 1a). Autoradiography of parallel samples readily detected CK2-dependent phosphorylation of the N-terminal WRN fragment and mass spectrometry analysis of the unlabelled *in vitro* phosphorylated N-terminal fragment revealed modification by CK2 within the acidic domain of WRN at multiple sites, including the six putative ones. An additional uncertain identification involved modification at S426, which has been recently found as a CDK2 substrate (*21*) (Supplementary Figure 1b). Mass spectrometry of the full-length WRN protein transiently expressed in HEK293T cells confirmed the identification of the six putative CK2-targeted S/T residues, either in the absence of aphidicolin treatment or upon replication stress (Supplementary Figure 2a, b). Of note, the identified residues were highly conserved in vertebrates (Supplementary Figure 2c).

**Fig. 1.**
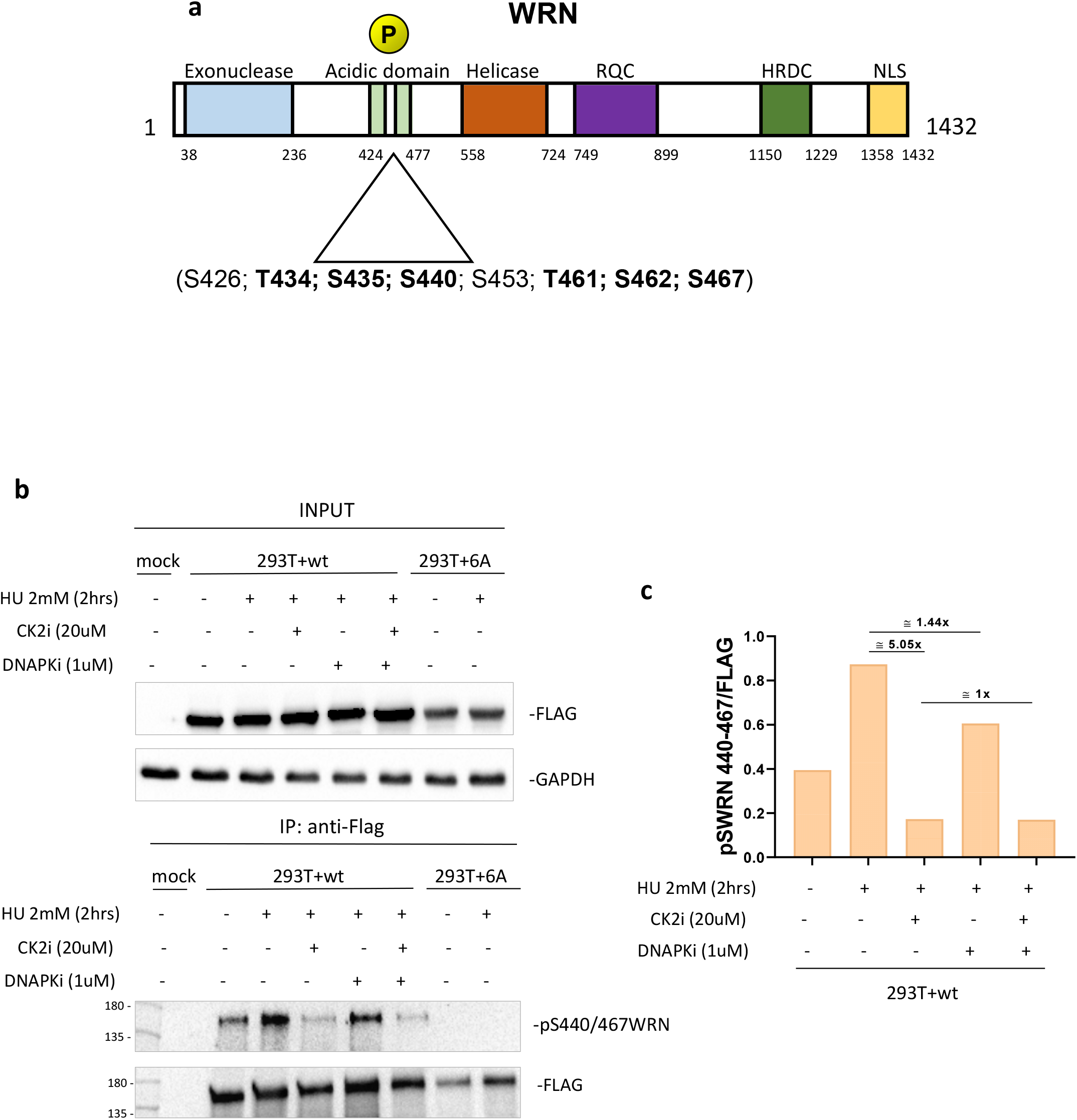
The acidic domain of WRN is phosphorylated by CK2. A) Schematic representation of WRN protein and domains. Mutation of six putative CK2 phosphorylation sites in the WRN acidic domain are highlighted. B) Anti-Flag-immunoprecipitates from HEK293T cells transfected with Flag-WRN or Flag-WRN^6A^ constructs. Cells were treated as indicated 48h after transfection. C) Quantification of WRN S440-467 phosphorylation sites and effect of CK2 or DNA-PK inhibition.

Some of the WRN residues identified as CK2 targets in our study, have also been reported to be modified by DNA-PK after DNA damage (*22*). To demonstrate that the acidic domain of WRN could be targeted by CK2 in response to replication arrest, we generated a phospho-specific antibody against WRN phosphorylated at S440 and S467 (pS440/467WRN). This antibody was used to probe cell lysate samples by Western blotting obtained after immunoprecipitation of Flag-tagged WRN wild-type or 6A transiently expressed in HEK293T cells. Forty-eight hours after transfection, cells were treated with HU for 2h in the presence of the CK2 inhibitor CX-4945 (CK2i) or the DNA-PKcs inhibitor NU7441 (DNA-PKi) to assess contribution of the two kinases to WRN phosphorylation. Analysis of anti-Flag IP with the anti-pS440/467WRN revealed that phosphorylation was already detectable in the absence of treatment but increased substantially in response to HU-induced replication arrest (Figure 1b, c). Noteworthy, no anti-pS440/467WRN signal was detectable by Western blotting from HEK293T cells transfected with the unphosphorylable WRN 6A mutant, supporting specificity of the antibody for the two residues (Figure 1b). Inhibition of CK2 was sufficient to decrease HU-induced phosphorylation approximately 5-fold, whereas inhibition of DNA-PK reduced WRN phosphorylation by only 1.4-fold. Combination of the two kinase inhibitors had no further effect as compared with treatment by CK2i alone (Figure 1c).

Collectively, these results demonstrate that the acidic domain of WRN is phosphorylated at multiple residues by CK2 *in vitro* and in human cells, and that CK2 but not DNA-PK, which was previously shown to target the WRN acidic domain (*22*), is the primary kinase that engages WRN in response to replication stress.

### Phosphorylation of the WRN acidic domain by CK2 drives its association with RPA

Having demonstrated that the WRN acidic domain is targeted by CK2 at six different sites *in vitro* and in cells, we sought to determine if this CK2-dependent phosphorylation contributes to the association of WRN with RPA. Thus, we generated a WRN fragment containing only the acidic domain (aa403-503), expressed it in bacteria as GST-fusion and used this purified fragment as bait in pull-down assays after *in vitro* phosphorylation with CK2 (Figure 2a). As a control, we expressed the corresponding fragment with the six S/T>D mutations mimicking the phosphorylated status (Figure 2a). The presence of RPA32 was used as readout of the interaction with the RPA heterotrimer, and the phosphorylation status of the CK2 sites was inferred using the anti-pS440/467WRN antibody. The wild-type fragment of WRN was greatly phosphorylated as shown by the anti-pS440/467WRN antibody staining, and a very minor cross-reactivity was detected in the phosphomimetic mutant fragment (Figure 2b). Consistent with a previous work (*17*), the mock-phosphorylated WRN^403-503^ fragment pulled-down RPA, but the amount of RPA32 bound to the fragment was increased more than 4-fold via prior phosphorylation by CK2 (Figure 2b). Of note, the phosphomimetic WRN6D^403-503^ fragment mimicked the increased association shown by the phosphorylated wild-type WRN fragment (Figure 2b). Phosphorylation status of the six S/T CK2-targeted residues of WRN also modulated its interaction with RPA in the cell. Indeed, Co-IP experiments, performed with extracts from HEK293T cells transiently transfected with Flag-tagged wild-type or unphosphorylable S/T>A (6A) WRN protein, showed that association of WRN with RPA increased during replication fork arrest induced by HU and that this association was strongly reduced for the unphosphorylable mutant (Figure 2c).

**Fig. 2.**
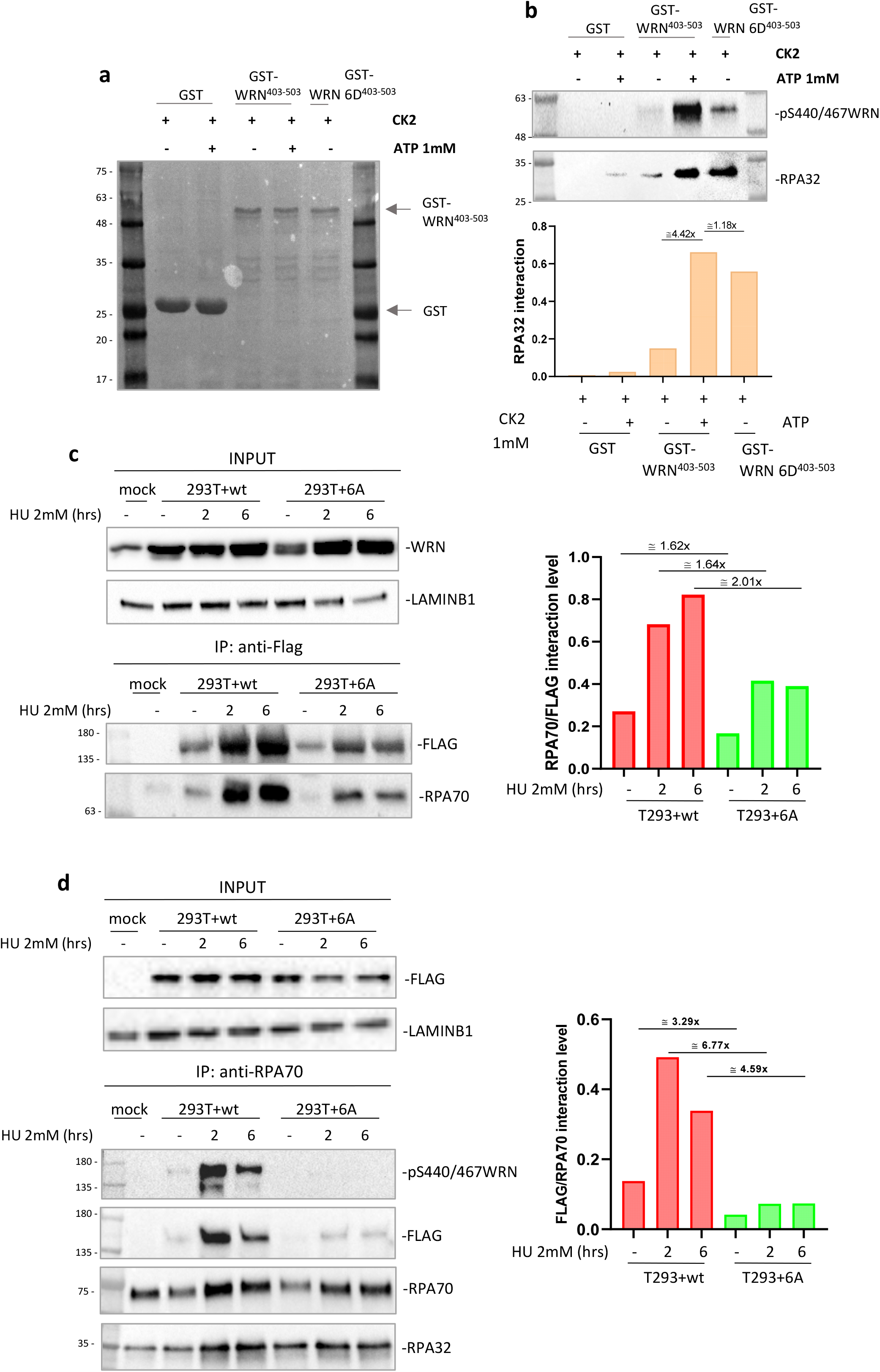
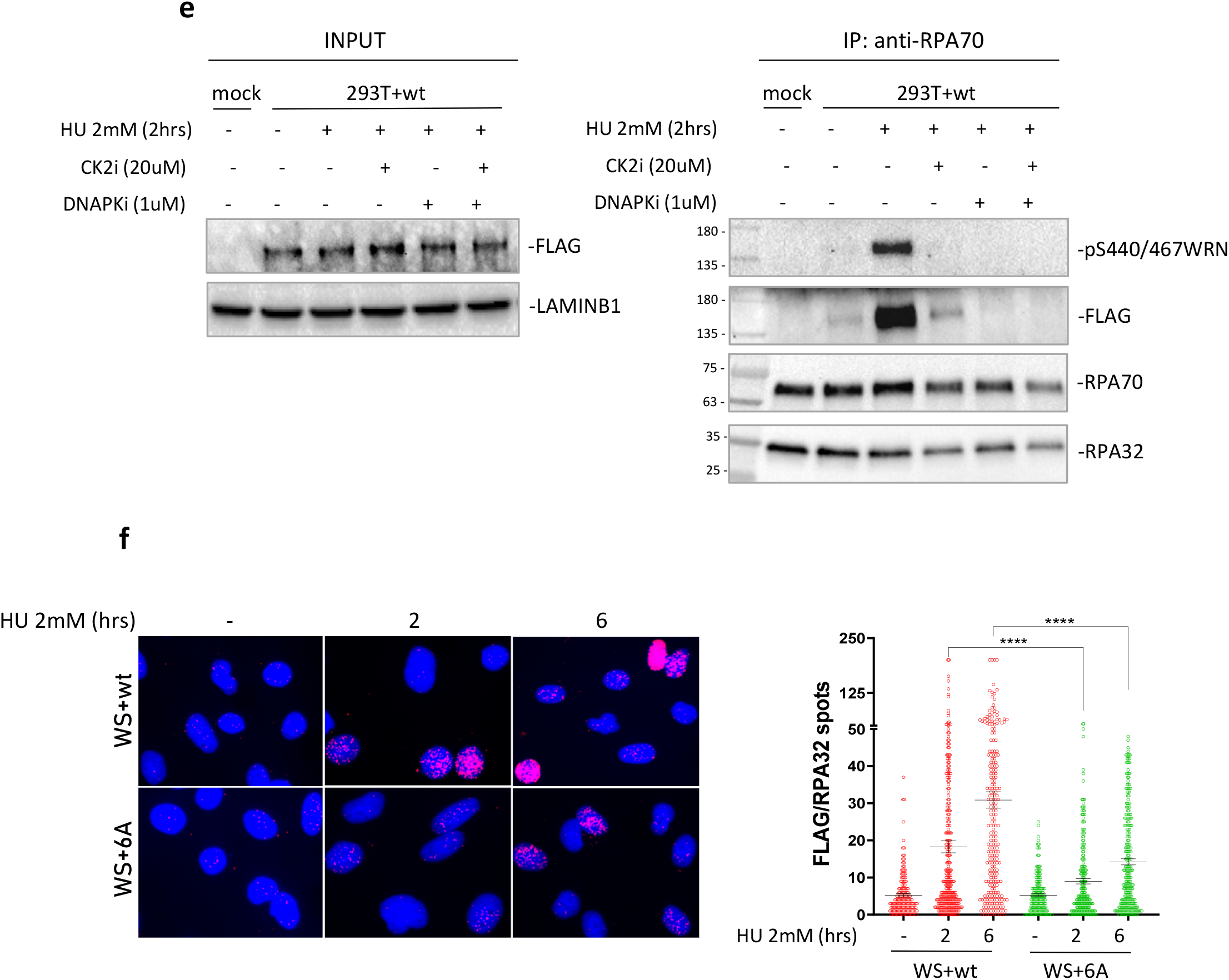
Phosphorylation of the acidic domain of WRN by CK2 drives association with RPA. A) Ponceau staining of GST-pulldowns with HEK293T nuclear extracts and GST-tagged WRN fragment 403-503 (WRN^wt^ and WRN^6D^) previously phosphorylated by CK2 in the presence or not of ATP. B) Western Blotting analysis of WRN S440/467 phosphorylation and RPA32 subunit from GST-pulldowns. The graph shows the levels of S440/467 phosphorylation and RPA32 bound to GST-tagged WRN fragments. C) Anti-Flag-immunoprecipitation from HEK293T cells transfected with Flag-WRN or Flag-WRN6A constructs. The graph shows the quantification of the WRN-normalised amount of RPA70 in the anti-Flag immunoprecipitate from the representative experiment. D) Anti-RPA70 immunoprecipitatation from HEK293T cells transfected with Flag-WRN or Flag-WRN6A constructs. The fraction of pS440/467 phosphorylated WRN associated with RPA was analysed using the anti-pS440/467WRN antibody. The graph shows the quantification of the RPA70-normalised amount of WRN in the anti-RPA70 immunoprecipitate from the representative experiment. E) Anti-RPA70 immunoprecipitation from HEK293T cells transfected with Flag-WRN^wt^ construct. Cells were treated with HU 2mM for 2h in the presence or not of the indicated inhibitors. F) WRN/RPA32 interaction was detected in Werner Syndrome (WS) cells nucleofected with Flag-WRN^wt^ or Flag-WRN^6A^ by in situ Proximity Ligation Assay using anti-FLAG and RPA32 antibodies. The graphs show individual values of PLA spots. Representative images are shown. Bars represent mean ± S.E. (*P<0.05; **P< 0.01; ***P< 0.001; ****P< 0.0001. Where not indicated, values are not significant).

To further demonstrate the relevance of phosphorylation status of the WRN acidic domain for its association with RPA, we immunoprecipitated RPA70 from cells transiently expressing WRN wild-type or 6A and analysed the presence of WRN and its phosphorylation status by Western blotting. As shown in Figure 2d, the interaction of RPA with WRN was enhanced already at 2h of HU exposure and remained high at 6h. Of note, the level of S440/467 phosphorylation followed the same trend, suggesting that most, if not all, of WRN residing in the complex with RPA is modified by CK2. Consistent with this, extremely low levels of WRN were found co-precipitating with RPA from cells expressing the unphosphorylable WRN 6A mutant, as assessed by anti-Flag Western blotting (Figure 2d).

To further assess if WRN requires phosphorylation by CK2 to associate with RPA, we immunoprecipitated RPA70 from cells transiently expressing WRN wild-type and treated with HU in the presence or not of the CK2i, DNA-PKi or both. As expected, co-immunoprecipitation of RPA and WRN was stimulated by replication arrest and was abrogated almost completely by inhibition of CK2 (Figure 2e). However, inhibition of DNA-PK also prevented formation of the WRN-RPA complex after replication arrest (Figure 2e). Since phosphorylation of WRN was largely dependent on CK2 after 2h of HU exposure (Figure 1b, c), we surmise that DNA-PK may control the interaction of RPA with WRN by targeting different WRN residues outside the acidic domain or, indirectly, by targeting RPA or other proteins.

To demonstrate that the interaction of WRN and RPA is dependent on the phosphorylation status of the WRN acidic domain at the single-cell level and by an orthogonal assay, we performed anti-Flag/RPA32 PLA in WS-derived patient cells complemented with the Flag-tagged wild-type WRN protein or its 6A unphosphorylable mutant. Consistent with co-IP data, PLA showed that interaction of WRN with RPA is strongly stimulated by replication arrest in a time-dependent manner and that it was suppressed in the absence of phosphorylation (Figure 2f).

Since association with RPA is often required to help recruit proteins to blocked replication forks, we performed PLA experiments to monitor the association of WRN 6A with parental ssDNA, an intermediate that accumulates at blocked replication forks, or nascent ssDNA, an intermediate that is formed at reversed forks or at processed collapsed forks. However, despite its extremely impaired ability to associate with RPA, the WRN 6A mutant retained almost complete proficiency to bind to ssDNA exposed at parental or nascent strand after replication arrest (Supplementary Figure 3a, b). Consistent with this evidence, chromatin localisation of the WRN wild-type and 6A was only slightly reduced after HU exposure (Supplementary Figure 3c).

Altogether, these results indicate that the association of WRN with RPA is strongly dependent on the phosphorylation status of six CK2-targeted residues in the acidic domain of WRN and that its interaction with RPA is only partially required for WRN recruitment in response to replication fork arrest.

### Interaction of WRN with RPA is not required for end-resection at blocked or collapsed replication forks or to prevent DSB formation during replication stress

In our hands, deletion of the WRN acidic domain led to extremely reduced expression of the protein, perhaps due to its destabilisation in the cell (Supplementary Figure 4a, b). Thus, the unphosphorylable WRN 6A mutant, which displayed a compromised ability to bind RPA, but normal expression and association with perturbed forks, can be useful to probe the functional role of the WRN-RPA interaction in the cell. At perturbed replication forks, WRN has been shown to play critical functions, assisting DNA2 during the physiological exonucleolytic processing at reversed forks and limiting engagement of pathological degradation by MRE11 (*23*, *24*). Contribution of the WRN-RPA interaction in both these mechanisms is unknown. As a proxy for the degradation occurring at reversed forks (*24*), we first evaluated accumulation of nascent ssDNA at different times of treatment with HU in WS cells complemented with the wild-type or the 6A WRN protein. Exposure of nascent ssDNA increased significantly after 6h of HU exposure for cells expressing the wild-type WRN and similarly in cells expressing the RPA-binding defective 6A mutant (Figure 3a). To further assess that RPA-binding of WRN was not involved in this function, we performed the DNA fiber assay (Figure 3b). Nascent DNA was sequentially pulse-labelled with CldU and IdU followed by treatment with HU. Analysis of IdU/CldU ratio revealed no statistically significant difference between WS cells complemented with wild-type WRN or its 6A mutant (Figure 3b). However, in both conditions, the IdU/CldU ratios were increased by the MRE11i, MIRIN, confirming that a fraction of forks underwent degradation after 6h of HU exposure, irrespective of the WRN binding to RPA.

**Fig. 3.**
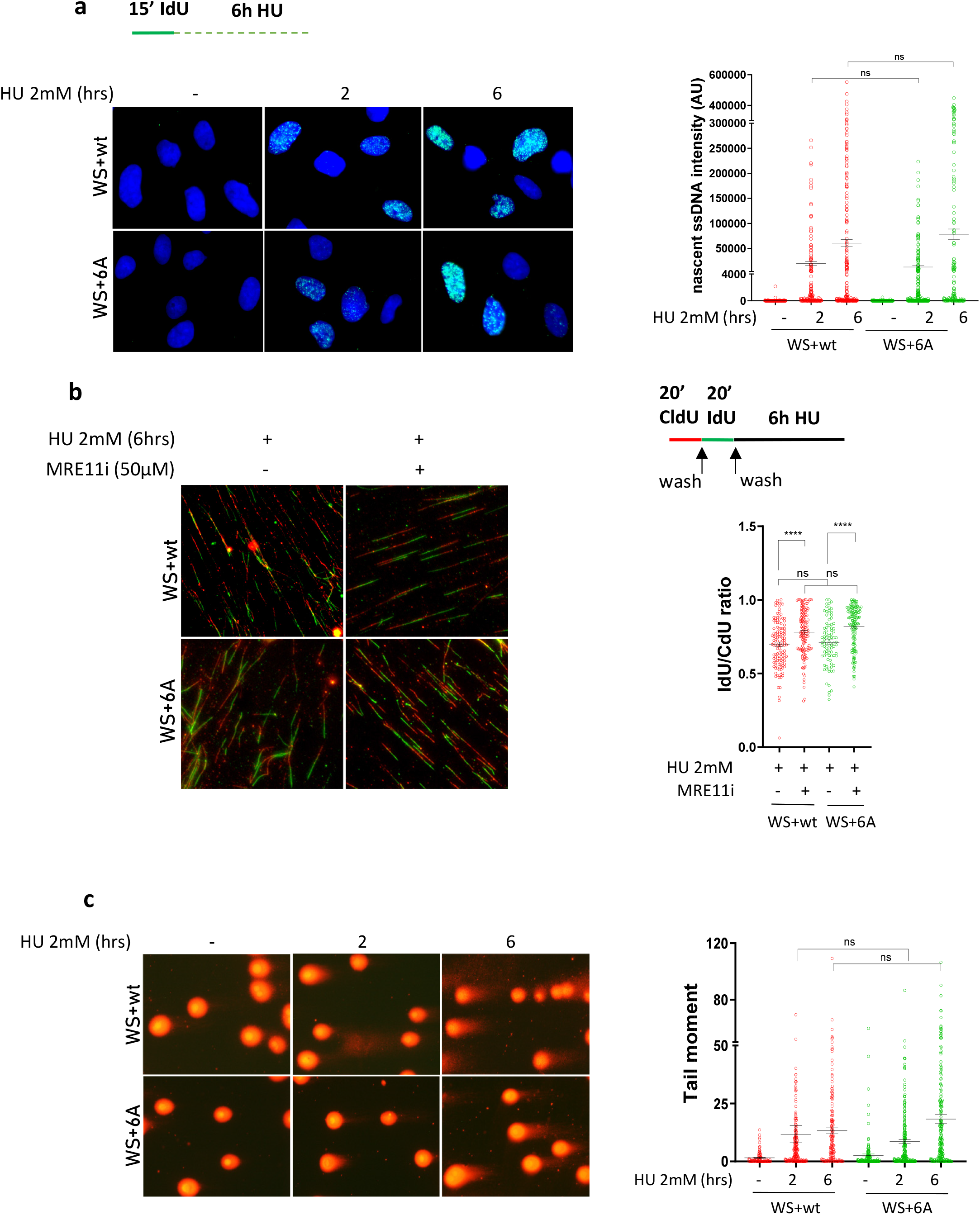
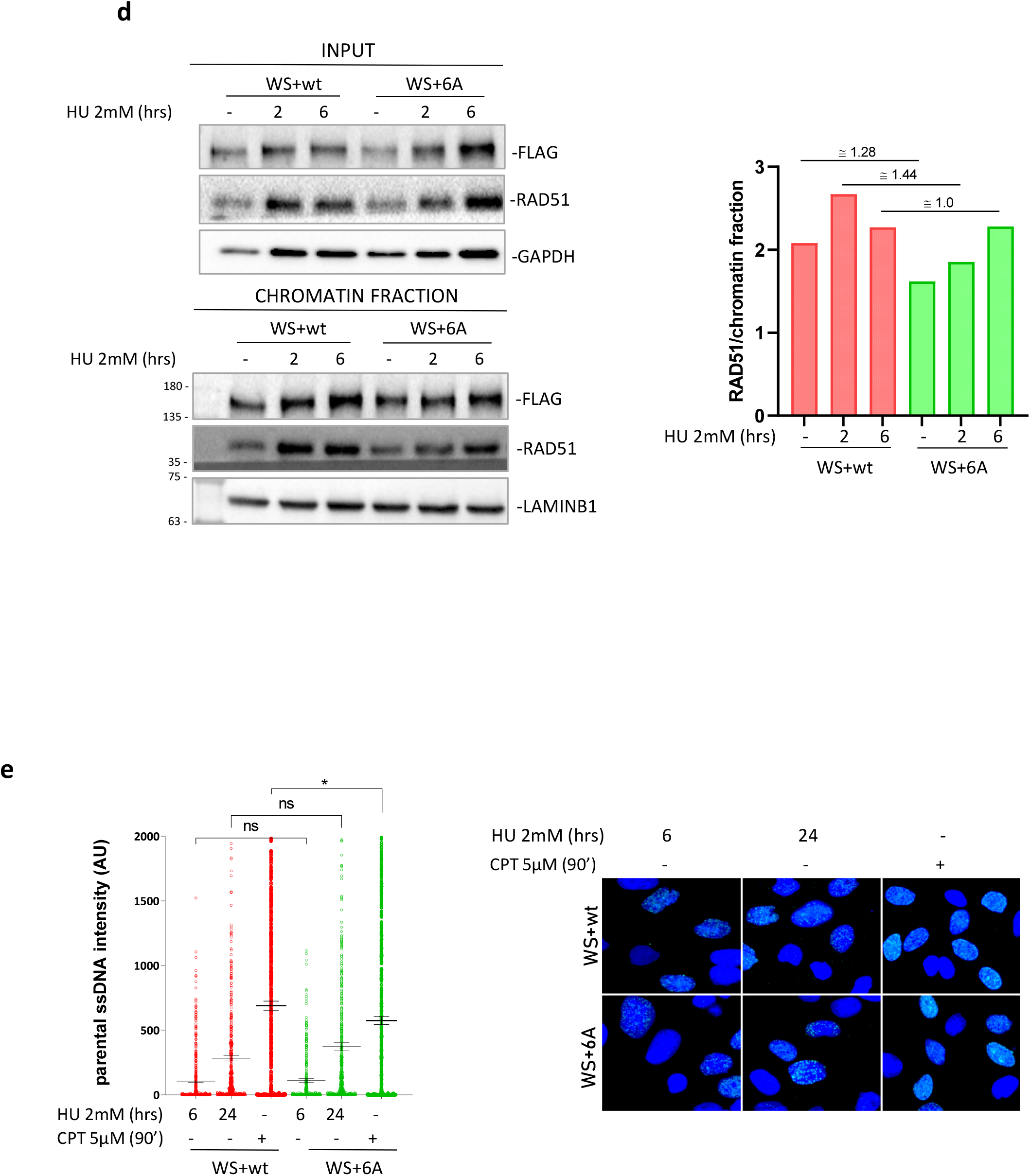
Interaction of WRN with RPA is not required for end-resection at blocked or collapsed replication forks or to prevent DSBs formation during replication stress. A) Detection of nascent ssDNA by immunofluorescence in WRN^wt^ and WRN^6A^ nucleofected WS cells treated as in the scheme. The graph shows individual values of IdU foci intensity (n=3). Bars represent mean ± S.E. (ns = not significant). Representative images are shown in the panel. B) Analysis of IdU/CdU ratio using DNA fiber assay (experimental scheme on top). The graph shows the individual IdU/CdU ratio values from duplicate experiments. Bars represent mean ± S.E. (ns = not significant; ****P< 0.0001.) Representative images of DNA fibers from random field are shown in the panel. C) Analysis of DSBs by neutral Comet assay. WRN^wt^ and WRN^6A^ nucleofected WS cells were treated as indicated. The graph shows individual tail moment values from two independent experiments. Bars represent mean ± S.E. (ns = not significant). D) Analysis of recruitment in chromatin of WRN and RAD51. Chromatin fractions were analysed by Western Blot. Input represents 1/40 of the chromatin fractionated lysate. The graph shows the quantification of the RAD51 amount in chromatin, normalised against LAMINB1 from the representative experiment. E) Analysis of end-resection by native IdU/ssDNA assay. WRN^wt^ and WRN^6A^ nucleofected WS cells were treated as indicated. Graph shows the quantification of total IdU intensity in 300 nuclei from three-independent experiments. Bars represent mean ± S.E. (ns = not significant; *P<0.05).

Loss of WRN function triggers formation of DSBs and stimulates RAD51-dependent repair (*25*, *26*). Thus, we performed neutral Comet assays to evaluate DSBs after replication arrest at various times of HU. As expected, some DSBs were found at 6h of HU treatment in WS cells complemented with the wild-type WRN protein, however, no difference was observed in the cells expressing the 6A mutant (Figure 3c). Consistent with the absence of any increased formation of DSBs after replication arrest for cells expressing the RPA-binding defective WRN mutant, chromatin fractionation experiments did not reveal a differential recruitment of RAD51 to chromatin in these cells (Figure 3d).

WRN has been implicated in promoting long-range end-degradation at collapsed replication forks (*27*, *28*). To test if loss of WRN-RPA binding affected end-resection at collapsed forks, we measured the formation of ssDNA after treatment with the topoisomerase inhibitor camptothecin (CPT), which induces replication stress and DSBs, by the native IdU/ssDNA assay (*28*). Formation of ssDNA as a readout of end-resection at CPT-induced DSBs was carried out in WS cells complemented with the WRN wild-type, the WRN 6A mutant or the end-resection defective S1133A WRN mutant (*28*). As shown in Figure 3e, WS cells complemented with wild-type WRN readily accumulate ssDNA in response to CPT treatment or 24h HU, another condition resulting in DSBs formation at replication forks. Expression of the RPA-binding defective 6A WRN mutant did not affect WRN’s activity during end-processing of DSBs formed at replication forks because the level of ssDNA detected in cells expressing the WRN 6A mutant was comparable with that observed in wild-type cells (Figure 3e).

Collectively, these results suggest that association of WRN with RPA is not implicated in the processing of stalled/collapsed forks, or in the prevention of DSBs.

### RPA-binding is required for WRN to restart replication fork and recover from replication arrest

Given that RPA-binding by WRN is dispensable during end-processing at perturbed replication forks, we next analysed if the WRN-RPA interaction affected replication fork restart and recovery. To this end, WS cells transiently complemented with the wild-type or the WRN 6A mutant, were pulse-labelled with CldU and treated with HU for 6h followed by 20min recovery in IdU before the spreading of DNA fibers (see scheme in Figure 4a). The analysis of the IdU/CldU ratios from DNA fibers showed shorter IdU tracts in cells expressing the RPA-binding deficient WRN 6A mutant (Figure 4a). To examine this further, we analysed replication fork dynamics in shWRN HEK293T transiently complemented with empty-vector, WRN wild-type or 6A (Supplementary Figure 5). Consistent with results in Figure 4a, analysis of the replication fork velocity revealed that RPA-binding is also important for WRN to support normal replication dynamics under conditions in which cellular replication is not challenged exogenously.

**Fig. 4.**
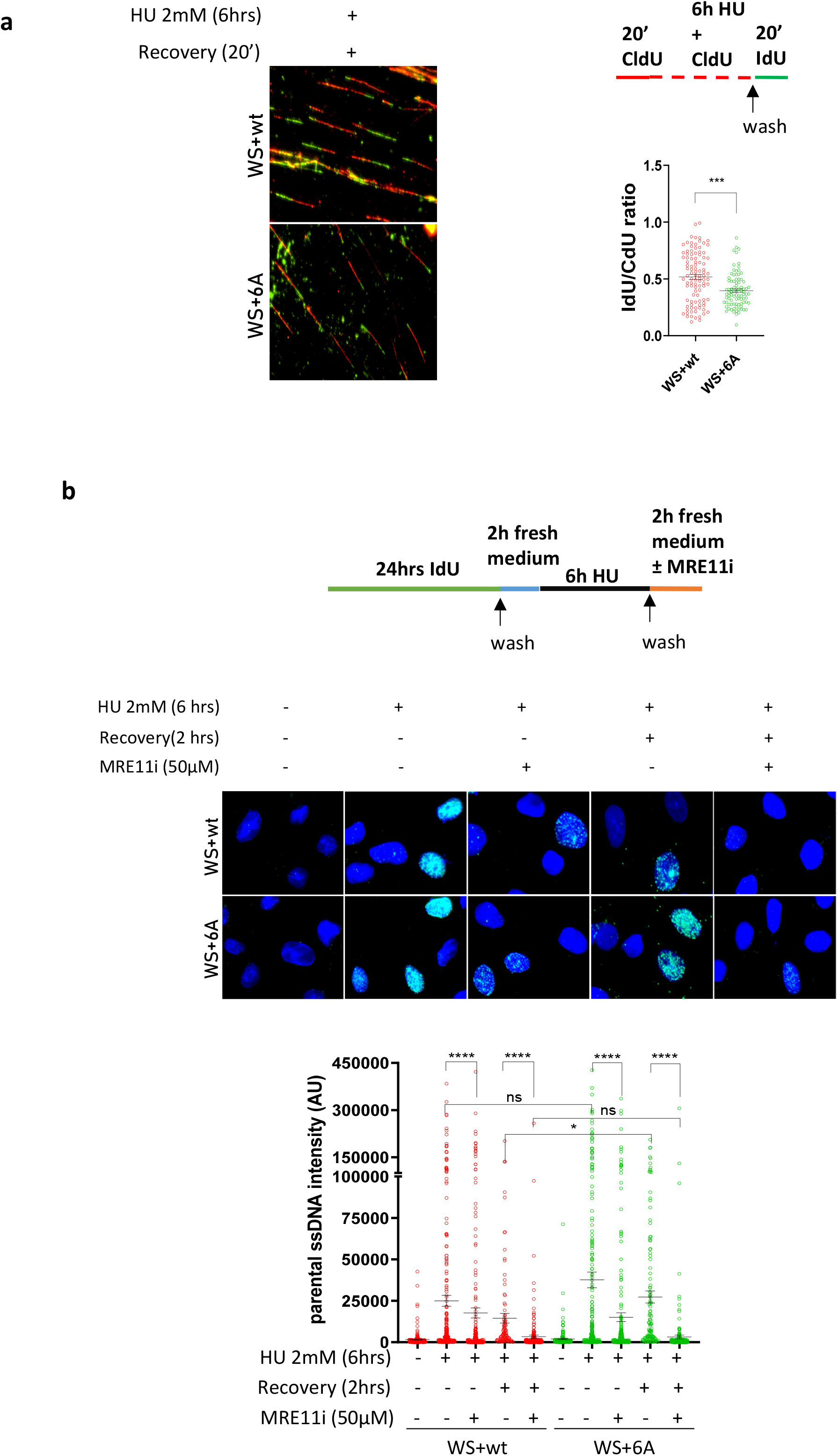
RPA-binding is required for WRN to restart replication fork and recover from replication arrest. A) Analysis of fork restart in IdU/CdU-labelled fibers from WRN^wt^ and WRN^6A^ expressing cells (Experimental scheme on top). The graph shows the individual IdU/CdU ratio values from duplicated experiments. Bars represent mean ± S.E. Representative images are shown. B) Analysis of parental ssDNA exposure in WRN^wt^ and WRN^6A^ expressing WS cells treated as indicated in the experimental scheme. The graph shows quantification of total IdU intensity for each nucleus from three independent experiments. Bars represent mean ± S.E. Representative images of native anti-IdU immunofluorescence are shown. (ns = not significant; *P<0.05; **P< 0.01; ***P< 0.001; ****P< 0.0001).

Since loss of RPA-binding impacts WRN’s ability to support DNA replication, we wondered whether cells expressing the WRN 6A mutant could show persistent ssDNA gaps after replication fork recovery. To test this possibility, we analysed the presence of parental ssDNA following recovery from 6h of HU by native IdU immunofluorescence in WS cells transiently complemented with the wild-type or 6A form of WRN (Figure 4b). Treatment with HU resulted in a fraction of cells showing some parental ssDNA exposure, but no significant difference was observed between WRN wild-type or 6A (Figure 4b). In both genetic backgrounds, a fraction of the parental ssDNA exposed during HU was dependent on MRE11 exonuclease activity because it was reduced by treatment with the MRE11i Mirin (Figure 4b). This was consistent with the DNA fiber degradation assay shown in Figure 3 and suggested that a sub-population of stalled forks were degraded after 6h of HU exposure in this cell line.

Of note, while the amount of parental ssDNA exposed in cells expressing wild-type WRN greatly decreased during recovery, this was not the case in cells expressing the WRN 6A mutant (Figure 4b). Strikingly, in both WRN wild-type or 6A, all the residual parental ssDNA exposed during recovery was MRE11-dependent, whereas DNA2-independent (Figure 4b and Supplementary Figure 6a). Inhibition of CK2 stimulated the exposure of parental ssDNA during HU treatment or recovery in wild-type cells, also exceeding the levels of parental ssDNA detected in the presence of WRN 6A (Supplementary Figure 6a). However, CK2 inhibition did not affect the amount of parental ssDNA observed in cells expressing the WRN 6A mutant (Supplementary Figure 6a).

These results suggest that in the absence of WRN-RPA binding cells accumulate parental ssDNA that is mostly dependent on MRE11. Since cells expressing WRN 6A did not show increased degradation at reversed forks when compared to wild-type cells, we wondered if the parental gaps accumulated because of repriming by PRIMPOL. To test this hypothesis, we repeated the analysis of the parental ssDNA in cells transfected or not with siRNA directed against PRIMPOL (Supplementary Fig. 6b). Based on other works, we expected that depletion of PRIMPOL would reduce exposure of parental ssDNA if gaps derived from its repriming activity. Interestingly, depletion of PRIMPOL failed to reduce the level of parental ssDNA exposed in cells expressing wild-type WRN as well as in cells expressing WRN 6A (Supplementary Fig. 6c).

Collectively, these results suggest that RPA-binding by WRN is important for accurate replication fork progression under unchallenged and, most importantly, perturbed conditions. Correct formation of the WRN-RPA complex allows cells to recover from perturbed replication without accumulating PRIMPOL-independent ssDNA gaps that become targets of MRE11-dependent degradation.

### RPA-binding by WRN collaborates with WRN helicase activity to promote proper replication and clearance of G4-DNA in cells

We show that RPA-binding by WRN is important for recovery of perturbed replication forks. In vitro, RPA facilitates WRN unwinding and fork regression making WRN a better helicase (*10*, *17*, *17*, *19*). Thus, we sought to determine if loss of RPA-binding and the helicase function of WRN acted in the same pathway by combining expression of WRN 6A and catalytic inhibition of the helicase using a small molecule inhibitor (WRNi; (*7*)). We first analysed the recovery of stalled forks using the DNA fiber assay (see scheme in Figure 5a). Pharmacological inhibition of WRN helicase activity strongly reduced the ability of cells expressing the wild-type form of WRN to recover stalled forks (Figure 5a). As expected, expression of the WRN 6A mutant that is defective in RPA-binding, reduced fork recovery (Figure 5a). Interestingly, although treatment of cells expressing WRN 6A with the WRNi reduced further the recovery of stalled forks, the effect was milder if compared to wild-type cells (Figure 5a). Of note, both the number of restarted forks and the fork progression during recovery (i.e., the length of the IdU tracts) were reduced by the compromised ability of WRN to bind RPA (Figure 5a).

**Fig. 5.**
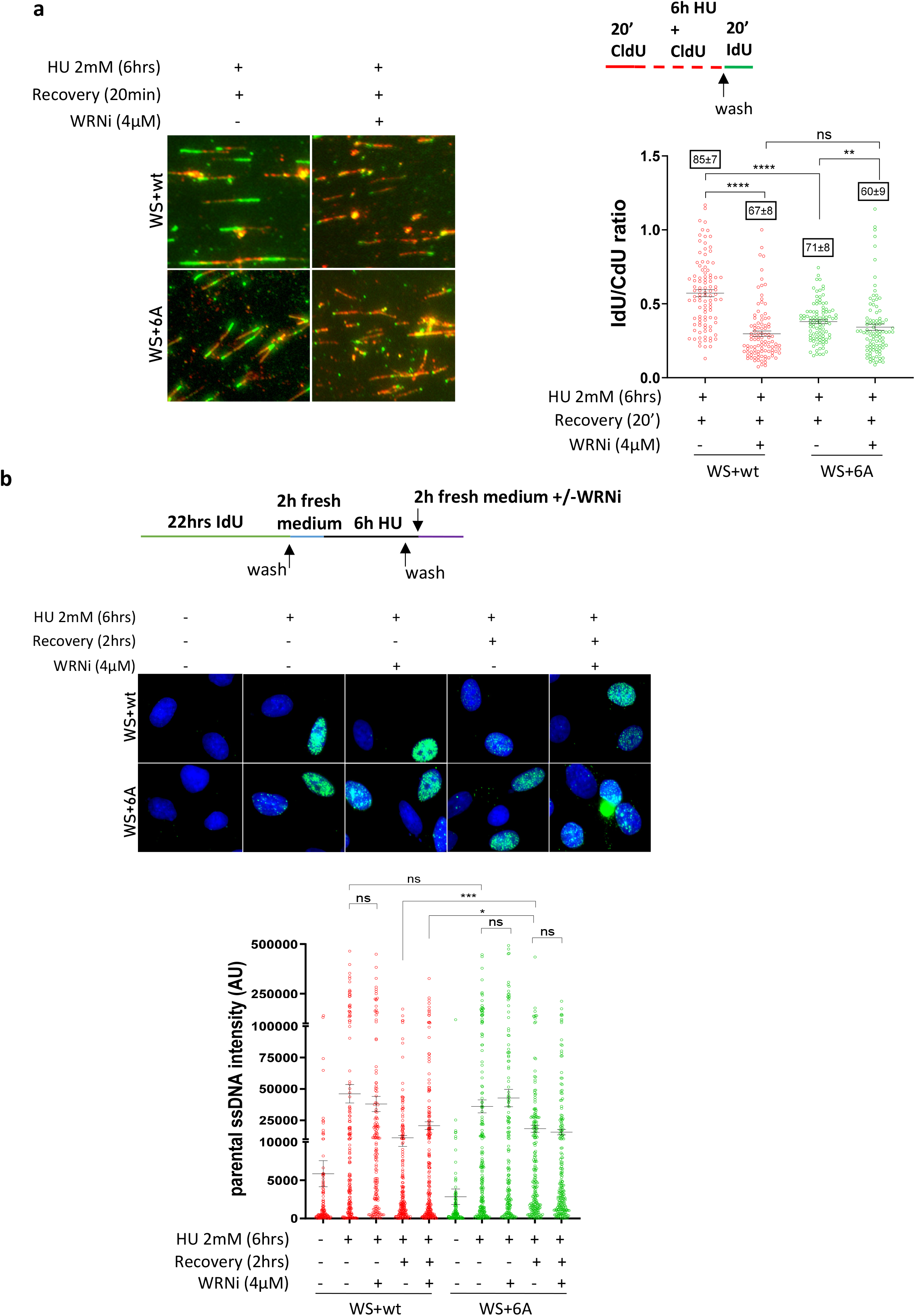
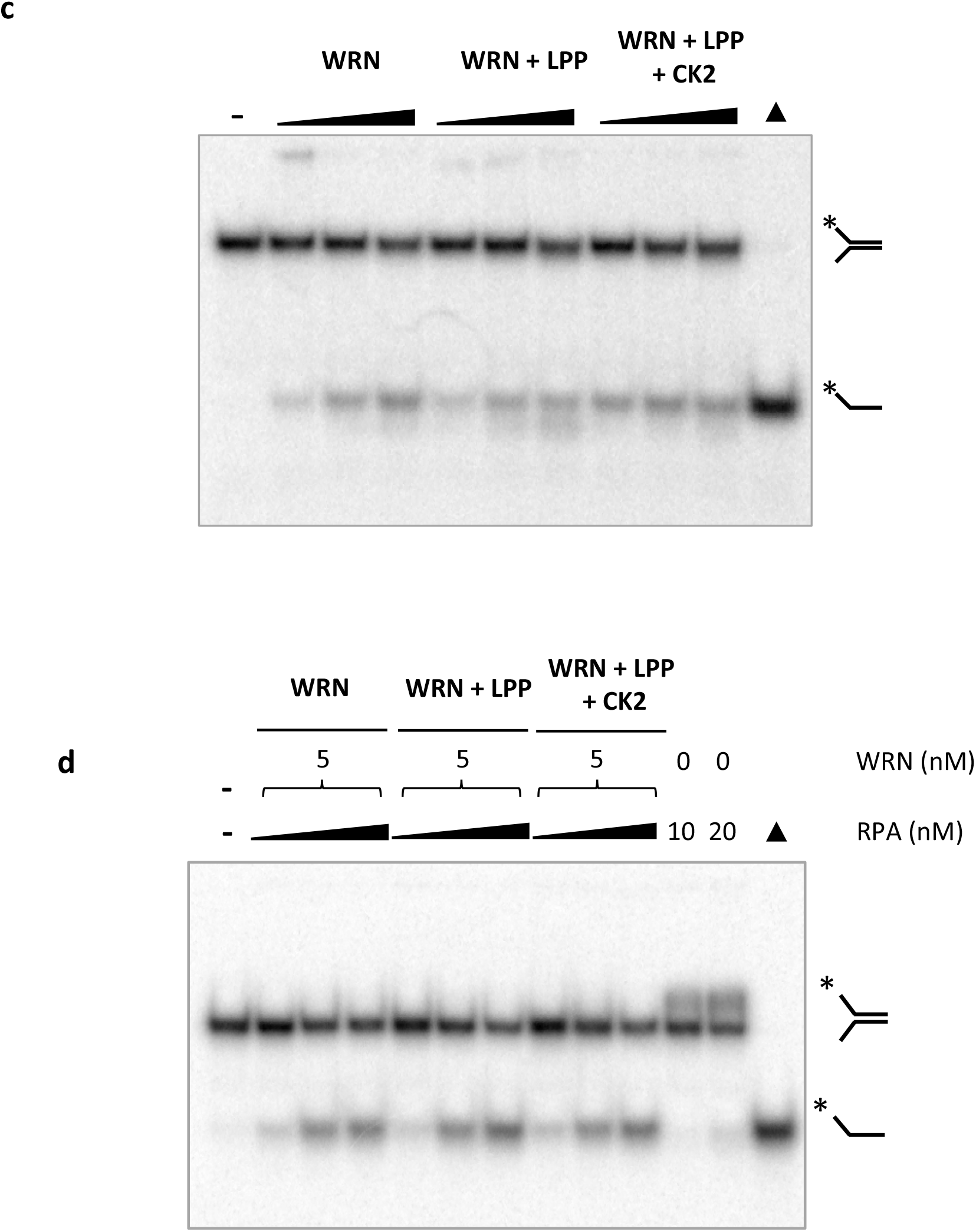
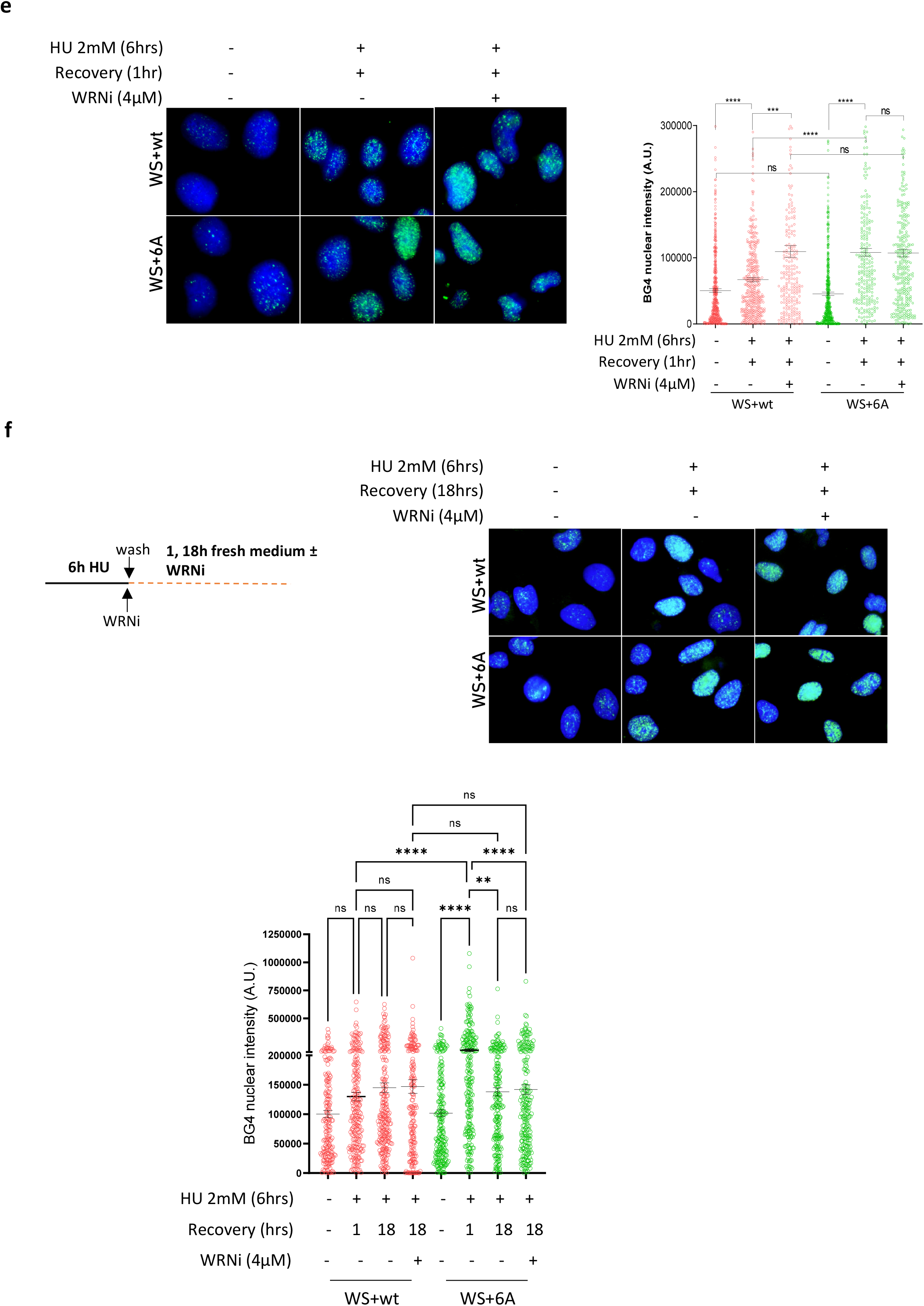
RPA-binding collaborates with WRN helicase activity of to promote replication recovery. A) Analysis of replication fork recovery from IdU/CdU-labelled DNA fibers as indicated in the experimental scheme in WS cells expressing Flag-WRN^wt^ or Flag-WRN^6A^. The graph shows the individual values of IdU/CdU ratio in isolated fibers from two independent replicates. Bars represent mean ± S.E. Representative images are shown. Numbers in the boxes above each dot plot are the % of restarting forks (mean ± S.E). B) Analysis of parental ssDNA exposure in WRN^wt^ and WRN^6A^ expressing WS cells treated as indicated. The graph shows quantification of total IdU intensity of each nucleus from three independent experiments. Bars represent mean ± S.E. Representative images are shown in the panel. C) Helicase activity of CK2-phosphorylated WRN. Helicase reactions containing 1.25, 2.5, 5 nM of untreated, LPP dephosphorylated, and LPP dephosphorylated-CK2 phosphorylated WRN were incubated with 25 bp forked DNA duplex substrate. D) Helicase activity of CK2-phosphorylated WRN in the presence of purified RPA heterotrimer. Fork substrate (15 nt arms, 34 bp duplex). ▲ denotes heat-denatured substrates. E) Analysis of G4 detection by an anti-DNA G-quadruplex structures antibody (clone BG4) in WRN^wt^ and WRN^6A^ expressing WS cells. The graph shows quantification of total BG4 nuclear staining for each nucleus from duplicate independent experiments. Bars represent mean ± S.E. F) Analysis of G4 detection by an anti-DNA G-quadruplex structures antibody (clone BG4) in WRN^wt^ and WRN^6A^ expressing cells during recovery. The graph shows quantification of total BG4 nuclear staining for each nucleus from duplicate independent experiments. Bars represent mean ± S.E. (ns = not significant; *P<0.05; **P< 0.01; ***P< 0.001;****P< 0.0001).

Next, we investigated if the increased exposure of parental ssDNA observed during recovery from replication arrest in cells expressing WRN 6A might be phenocopied in WRN wild-type cells by treatment with the WRNi. As shown in Figure 5b, inhibition of WRN helicase resulted in more parental ssDNA being exposed in cells expressing WRN wild-type but not in cells expressing the WRN 6A mutant. Of note, wild-type cells treated with the WRNi exposed as much parental ssDNA as cells expressing the RPA-binding deficient WRN 6A mutant (Figure 5b).

Because inhibition of WRN helicase in wild-type cells mimicked the phenotype of the WRN 6A mutant, we performed *in vitro* assays to investigate if CK2 phosphorylation at the WRN acidic domain regulates its helicase activity. To this end, we purified recombinant wild-type WRN and, as a control, we verified if the recombinant WRN purified from insect cells might be already phosphorylated at the CK2 sites of the acidic domain by Western blotting using the anti-pS440/467 WRN antibody. Interestingly, the recombinant WRN wild-type purified from insect cells is phosphorylated at CK2 sites (Supplementary Fig. 7). Thus, we subjected the purified WRN to dephosphorylation/phosphorylation using lambda phosphatase and subsequent addition of CK2 kinase prior to analysing its enzymatic activity and comparing WRN’s activity to only the dephosphorylated form which was mock-treated after the dephosphorylation incubation. The dephosphorylated and CK2-rephosphorylated WRN proteins were tested for catalytic activity on a forked duplex DNA substrate that is unwound by WRN in the presence of ATP or degraded by WRN’s 3′–5′ exonuclease activity in the absence of ATP. As shown in Figure 5c,d, no apparent difference in the helicase or exonuclease activity was detected between the unphosphorylated WRN and CK2 rephosphorylated WRN recombinant proteins, also in the presence of RPA.

WRN has been shown to catalyse unwinding of G-quadruplex (G4) DNA substrates *in vitro*, and RPA binding can boost WRN activity (*20*, *29*). Thus, we sought to determine if the compromised ability to resume stalled forks by the RPA-binding deficient WRN was correlated with poor helicase activity towards G4s. To test this possibility, we first evaluated the presence of G4s by anti-BG4 immunofluorescence in WS cells expressing WRN wild-type or 6A, in the presence or absence of the WRNi. Untreated cells showed little BG4 staining irrespective of the RPA-binding capability of WRN (Figure 5e). During recovery from HU, cells expressing WRN wild-type displayed increased BG4 staining, which was further increased by the co-treatment of cells with the WRNi during HU exposure (Figure 5e). Of note, impaired ability of WRN to bind RPA substantially increased BG4 staining after recovery matching the values observed after inhibition of the WRN helicase in wild-type cells, and no further increase in BG4 staining was observed for cells expressing WRN 6A that were treated with the WRNi (Figure 5e).

Having shown that either impaired helicase activity of WRN or binding to RPA led to accumulation of G4s during restart of stalled replication forks, we wanted to assess if these persisting G4s were eventually resolved. To this end, WS cells complemented with WRN wild-type or the 6A mutant were treated with HU and recovered for 1 or 18h before evaluating the presence of G4s by BG4 immunofluorescence assay (see scheme in Figure 5f). To additionally investigate the contribution of the WRN helicase activity, parallel samples were treated with WRNi during the 18hrs recovery from HU. Interestingly, after 18 hrs of recovery, cells expressing WRN 6A or having the WRN helicase inhibited showed dissolution of G4s and returned to wild-type levels (Figure 5f).

Therefore, these results demonstrate that impairment of RPA-binding by WRN is sufficient to induce accumulation of G4s shortly after recovery from replication arrest mimicking the effect of WRN helicase inhibition, although CK2-dependent phosphorylation of WRN does not impair enzymatic activity on a forked duplex in vitro; furthermore, they show that G4s which accumulate when WRN’s binding to RPA or its helicase activity is impaired are resolved after prolonged recovery from HU.

### MRE11-dependent gaps and MUS81-dependent DSBs contribute to G4 clearance in the absence of WRN-RPA binding

Collectively, our data support a model in which the binding of WRN to RPA is necessary to recover replication forks and correctly replicate secondary DNA structures such as G4s. This prompted us to investigate the relationship between the persistent parental gaps generated by MRE11 and the removal of G4s observed at longer recovery times after replication fork arrest in cells expressing the WRN 6A.

Thus, we treated cells with HU and analysed the presence of G4s using immunofluorescence after 1 hour and 18 hours of recovery, with or without the MRE11 inhibitor Mirin. Inhibiting MRE11 exonuclease activity decreased BG4 staining in wild-type cells after 1 hour of recovery but had no effect after 18 hours (Figure 6a). This effect was significant, but the amount of HU-dependent G4s estimated by anti-BG4 immunofluorescence was exceptionally low in wild-type cells. In contrast, inhibiting MRE11 exonuclease activity greatly increased the already elevated anti-BG4 staining in cells expressing WRN 6A (Figure 6a). The observation that G4 removal depended on MRE11 exonuclease activity suggests that DSBs are formed and resected at G4 sites.

**Fig. 6.**
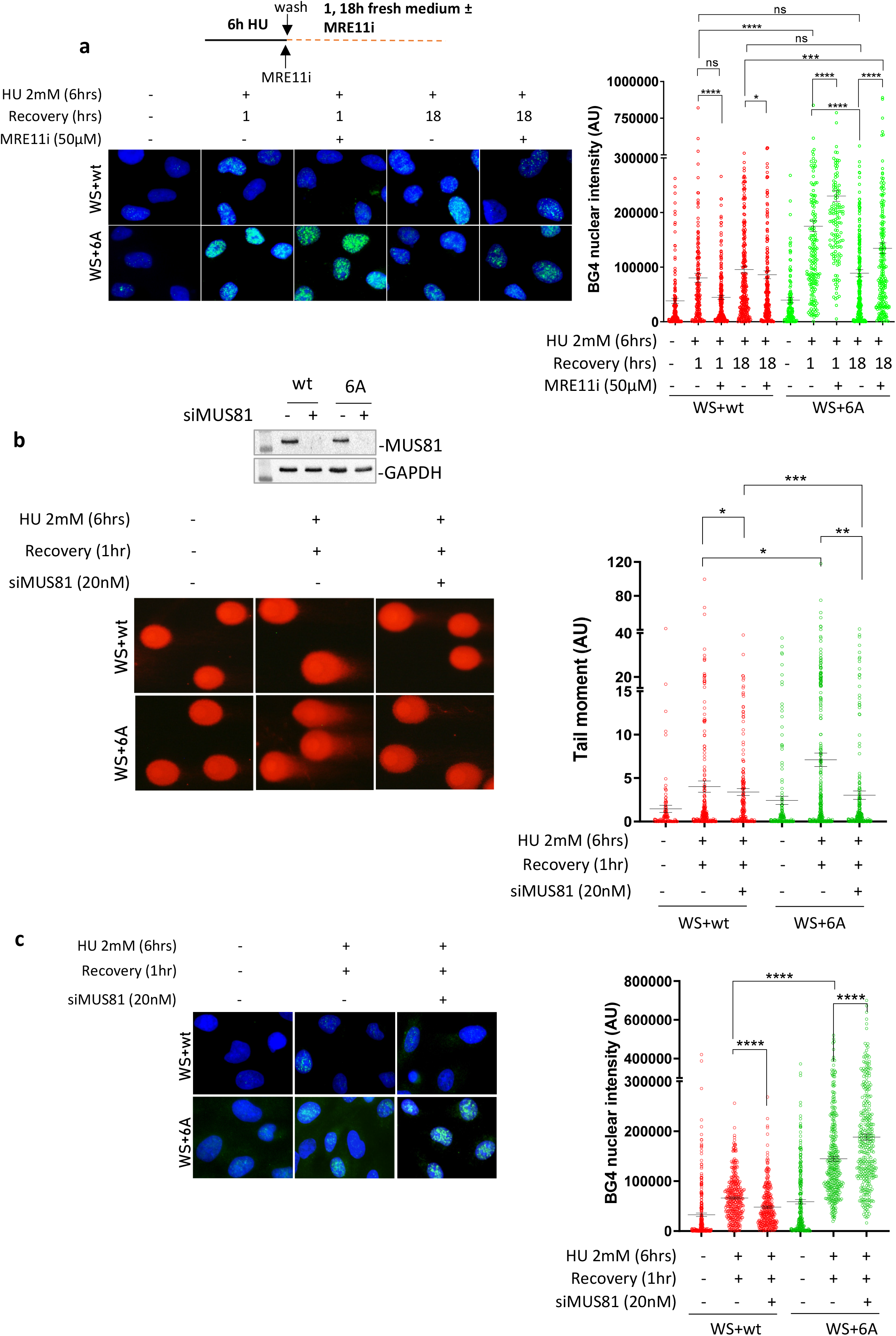
MRE11-dependent gaps and MUS81-dependent DSBs contribute to G4 clearance in cells that are defective for RPA binding to WRN. A) Analysis of G4s accumulation evaluated by anti-BG4 immunofluorescence in WRN^wt^ and WRN^6A^ nucleofected WS cells treated with HU and recovered in the presence or absence of the MRE11 exonuclease inhibitor Mirin (MRE11i). The graph shows the individual values of BG4 nuclear intensity. Bars represent mean ± S.E. (ns = not significant; *P<0.05; ***P< 0.001; ****P< 0.0001. Where not indicated, values are not significant). B) Analysis of DSBs by neutral Comet assay. WRN^wt^ and WRN^6A^ nucleofected WS cells were transfected with CTRL or MUS81 siRNA and 48h after treated with HU and recovered for 1 hour. WB shows the downregulation of MUS81. The graph shows individual tail moment values from duplicated experiments. Bars represent mean ± S.E. Statistical analyses were performed by Student’s t-test (*P<0.05; **P<0.01; ***P< 0.001. Where not indicated, values are not significant). C) G4s accumulation was detected by anti-BG4 immunofluorescence. Cells expressing the wild-type and mutant WRN were transfected with CTRL or MUS81 siRNA and, 48h after, they were treated with HU and recovered for 1 hour. The graph shows the individual values of BG4 foci nuclear intensity. Bars represent mean ± S.E. (****P< 0.001. Where not indicated, values are not significant).

In human cells, MUS81 endonuclease can process G4s and is known to introduce DSBs in the absence of WRN (*26*, *30*). We depleted MUS81 using RNAi (Figure 6b) and analysed whether DSBs formed during recovery from HU in cells expressing WRN 6A. The neutral comet assay showed very low levels of MUS81-independent DSBs in cells expressing wild-type WRN after 1 hour of recovery from HU (Figure 6b). However, many more DSBs were found in cells expressing WRN 6A during recovery, and these were completely suppressed by MUS81 depletion (Figure 6b). Of note, DSBs formed in cells expressing the WRN 6A mutant during recovery were also strongly reduced by Mirin (Supplementary Fig. 8), confirming that MRE11-dependent gap enlargement acts upstream of MUS81. This MUS81-dependent formation of DSBs led us to analyse the presence of G4s using anti-BG4 immunofluorescence to correlate them with G4 clearance (Figure 6c). Depletion of MUS81 reduced the already low level of BG4 staining in WS cells complemented with wild-type WRN after 1 hour of recovery. In contrast, depletion of MUS81 increased the level of G4s in cells expressing WRN 6A.

Altogether, these results provide evidence for a crucial role of MRE11 and MUS81 in the removal of G4 structures that fail to be resolved due to defective interaction of WRN with RPA during replication fork recovery; furthermore, these findings link gap processing with DSBs formation.

### RAD51 repair DSBs formed at G4 in the absence of WRN-RPA binding

Our findings demonstrate that resolving G4 structures without WRN-RPA involvement necessitates the presence of MRE11 and MUS81. MRE11-enlarged gaps can be utilized to attract RAD51 for post-replication gap repair. To investigate whether RAD51 is recruited to parental ssDNA exposed at DSBs formed by MUS81 and through MRE11-dependent degradation of the newly synthesized DNA, we conducted the parental ssDNA-protein PLA (*24*). As depicted in Figure 7a, RAD51 was found associated with parental ssDNA in cells expressing wild-type WRN after recovering from HU, and this specific association was minimally affected by inhibiting MRE11 exonuclease. Conversely, the expression of WRN 6A resulted in a higher level of RAD51 associated with parental ssDNA, which was significantly reduced by treatment with Mirin (Figure 7a). Furthermore, the recruitment of RAD51 to parental ssDNA was substantially, but not completely, diminished after depleting MUS81 in cells expressing WRN 6A (Figure 7b), suggesting that RAD51 may also participate in the repair of MUS81-dependent DSBs. We hypothesized that if RAD51 was engaged in post-replication repair, it would still be detectable at ssDNA during a later recovery period. Thus, we monitored the recruitment of RAD51 to parental ssDNA using PLA after 18 hrs of recovery. As illustrated in Figure 7c, RAD51 was observed to be associated with parental ssDNA in cells expressing the wild-type WRN at 18 hrs of recovery from HU, and this specific association was minimally affected by Mirin, which interferes with DSB formation (see Supplementary Fig 8). Conversely, the expression of WRN 6A, which stimulates MRE11 and MUS81-dependent DNA breakage, led to a higher amount of RAD51 associated with parental ssDNA, and this association was greatly suppressed by treatment with the MRE11 inhibitor Mirin (Figure 7c).

**Fig. 7.**
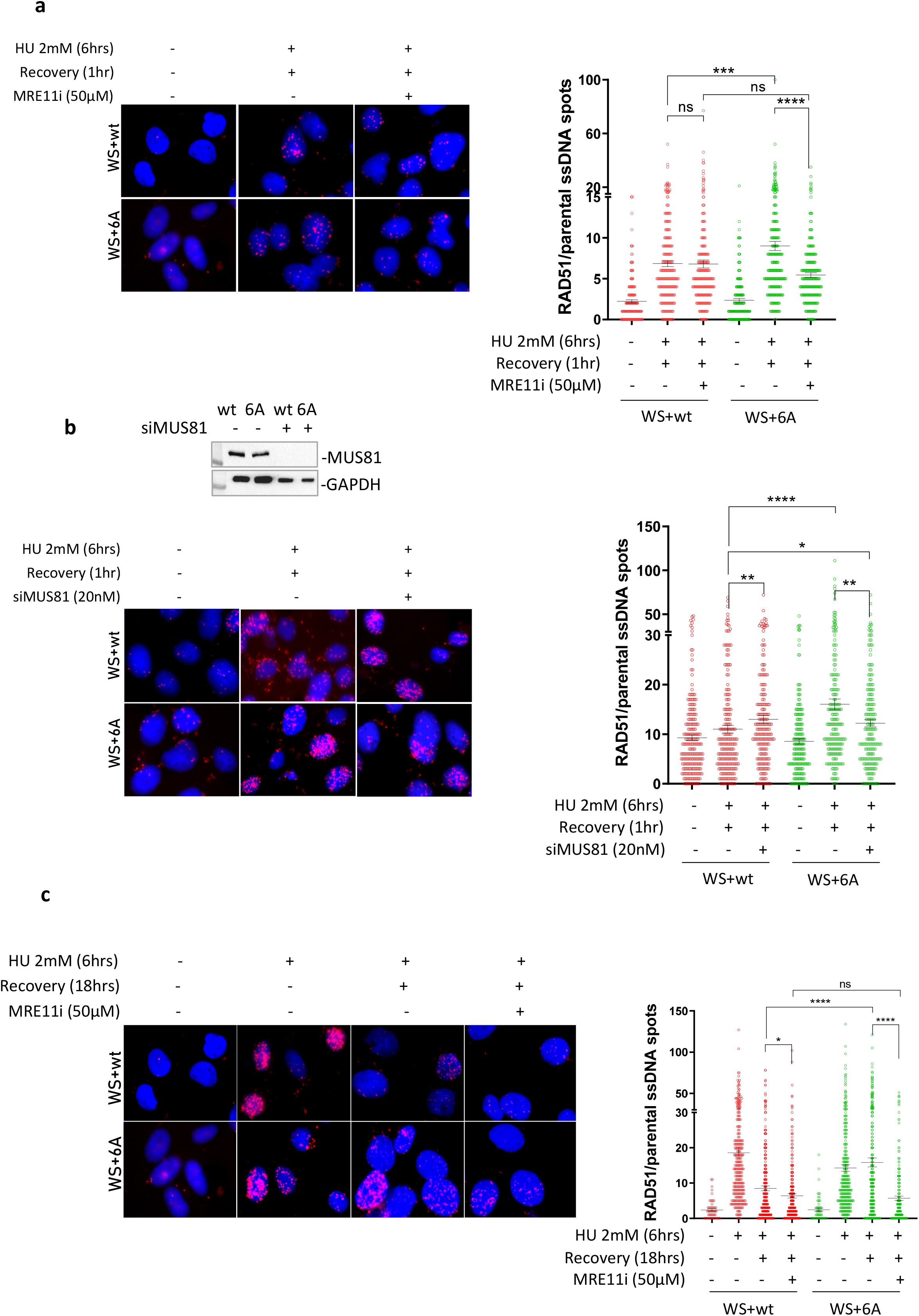
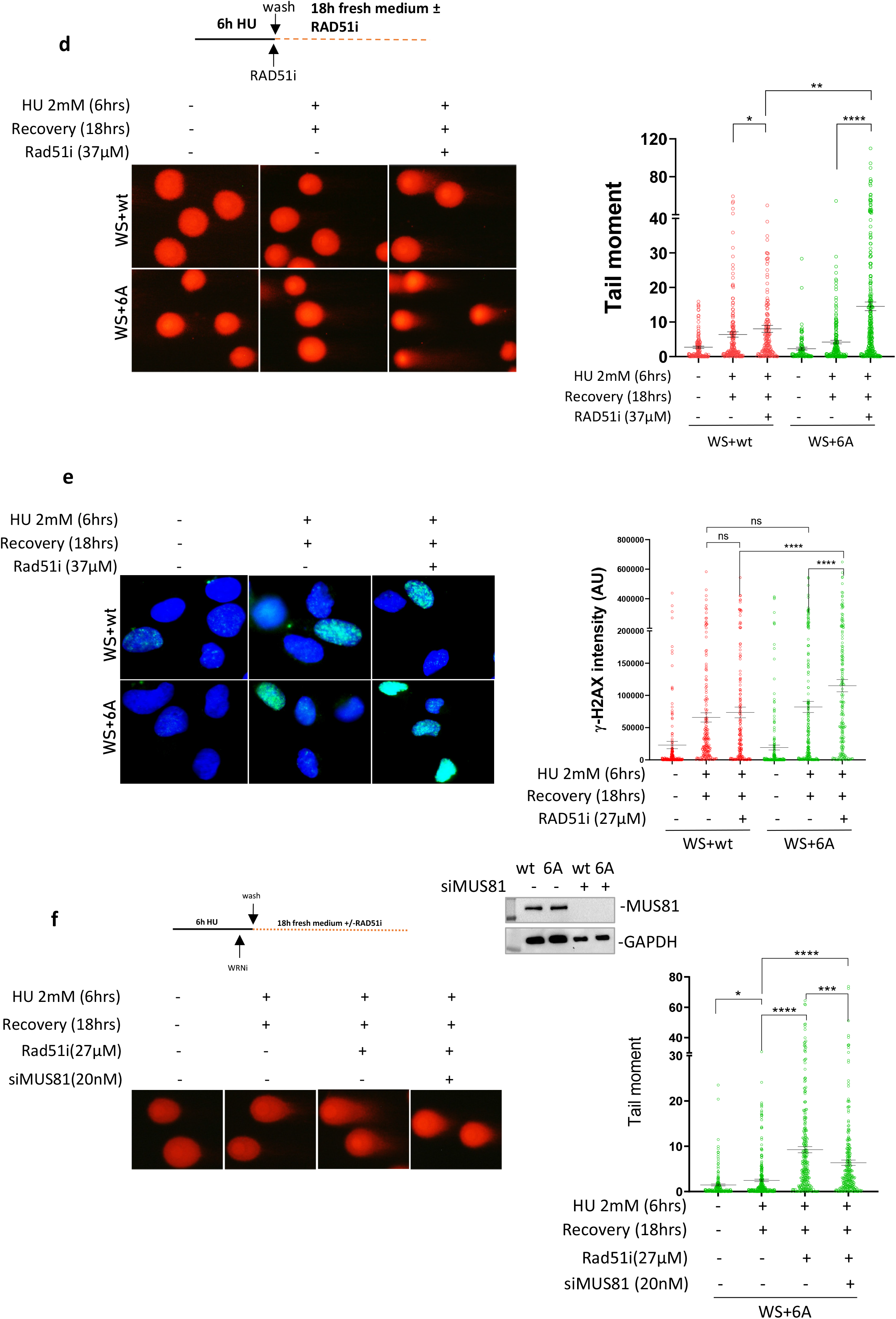
**RAD51 suppresses DSBs formed at G4 sites in the absence of WRN-RPA binding**. A) In Situ Proximity Ligation Assay between RAD51 and parental ssDNA. WS cells nucleofected with Flag-WRNwt or Flag-WRN6A were treated with HU and recovered for 1 hour in the presence or absence of the MRE11i Mirin. The graphs show individual values of PLA spots (n=2). Representative images are shown. Bars represent mean ± S.E. (ns = not significant; ***P<0.001; ****P< 0.0001. Where not indicated, values are not significant). B) In Situ Proximity Ligation Assay between RAD51 and parental ssDNA. WS cells nucleofected with Flag-WRNwt or Flag-WRN6A were transfected with CTRL or siMUS81 siRNA and treated 48h after with HU followed by recovery for 1 hour. WB shows the downregulation of MUS81. The graphs show individual values of PLA spots (n=2). Representative images are shown. Bars represent mean ± S.E. (ns = not significant; *P<0.05; **P<0.01; ****P< 0.0001. Where not indicated, values are not significant). C) In Situ Proximity Ligation Assay between RAD51 and parental ssDNA. WS cells nucleofected with Flag-WRNwt or Flag-WRN6A were treated with HU and recovered for 18 hours in the presence or absence of the MRE11i Mirin. The graphs show individual values of PLA spots (n=2). Representative images are shown. Bars represent mean ± S.E. (ns = not significant; *P<0.05; ****P< 0.0001). D) Neutral Comet assay for DSBs evaluation in WRNwt and WRN6A nucleofected cells during recovery from HU (see experimental scheme). The graph shows individual tail moment values (n=2). Bars represent mean ± S.E. Statistical analyses were performed by Student’s t-test (*P<0.05; **P< 0.01; ****P< 0.0001. Where not indicated, values are not significant). E) Analysis of anti-γ-H2AX immunofluorescence during recovery from HU (see experimental scheme). The graph shows the individual values of γ-H2AX foci intensity (n=2). Bars represent mean ± S.E. Statistical analyses were performed by ANOVA (ns = not significant; *P<0.05; **P< 0.01; ***P< 0.001; ****P< 0.0001). F) Analysis of DSBs by neutral Comet assay. WS cells nucleofected with Flag-WRN6A were transfected with CTRL or siMUS81 oligos and treated 48h thereafter with HU as indicated in the scheme. The graph shows individual tail moment values (n=2). Bars represent mean ± S.E. Statistical analyses were performed by Student’s t-test (*P<0.05; ***P< 0.001. ****P<0.0001).

Next, we reasoned that MUS81-dependent DSBs would persist even when RAD51 nucleofilament formation is blocked, if RAD51 is necessary for their repair. Therefore, we performed a neutral Comet assay on cells expressing the wild-type form of WRN or its RPA-binding-defective mutant at 18 hours of recovery, using the RAD51 inhibitor B02 (RAD51i). As shown in Figure 7d, only a few DSBs were present in WRN wild-type cells at 18 hrs of recovery from HU, and they were unaffected by RAD51 inhibition. Interestingly, cells expressing the RPA-binding-defective WRN also exhibited few DSBs at 18 hrs of recovery from HU, and there was no statistically significant difference compared to cells expressing WRN wild-type. However, when RAD51 was inhibited, the number of DSBs substantially increased (Figure 7d). Consistent with the neutral Comet assay, the phosphorylation level of the H2AX histone, which serves as a marker for DNA damage, was significantly elevated by RAD51 inhibition in cells expressing the RPA-binding-deficient WRN (Figure 7e).

Subsequently, we examined the persistence of DSBs after transfection with a control siRNA or an siRNA targeting MUS81 in cells expressing the RPA-binding-deficient WRN mutant at 18 hours of recovery in the presence of the RAD51 inhibitor. We hypothesized that despite RAD51 inhibition, DSBs would be reduced if MUS81 was silenced, indicating that RAD51 is primarily required for repairing MUS81-dependent DSBs. Analysis of the number of DSBs using the neutral Comet assay confirmed that RAD51 inhibition increased their count (Figure 7f). Furthermore, Figure 7f demonstrated that depletion of MUS81 substantially decreased the number of DSBs compared to control-depleted cells and RAD51-inhibited cells.

Altogether, these results suggest that RAD51 is recruited to MRE11-processed gaps and is necessary for repairing MUS81-dependent DSBs, thereby contributing to the clearance of G4 structures and limiting DNA damage.

## DISCUSSION

In this study, we determined that RPA binding to WRN plays a unique role at stressed replication forks in a manner that is dependent on post-translational phosphorylation of WRN that regulates its interaction with RPA. We identified a cluster of CK2-dependent phosphorylation sites in the acidic domain of WRN that are essential for its optimal interaction with RPA. Characterisation of a WRN 6A unphosphorylable mutant allowed us to pinpoint a biological function of the WRN-RPA interaction that is critical for genome stability.

Previous *in vitro* studies have shown that WRN interacts with RPA *via* its acidic domain that binds to a basic cleft in the N-terminal region of the RPA1 subunit (*10*, *17*, *19*). Consistent with these observations, we determine that phosphorylation of the WRN acidic domain by CK2 at multiple sites modulates the WRN-RPA interaction. Most importantly, our data indicate that, in the cell, most, if not all, of the WRN-RPA interactions are inhibited by abrogating phosphorylation at the acidic domain of WRN. As a minor RPA-binding site has been mapped to the C-terminal region of WRN (*17*), we cannot exclude that the very residual level of interaction observed in the WRN 6A mutant derives from this site.

Of note, the described CK2-dependent phosphorylation sites are evolutionary conserved, supporting their relevance. Indeed, they can be found also in vertebrate WRN (chicken) and in Xenopus FFA-1. Interestingly, two out of the six CK2-dependent phosphorylation sites identified in the present work, S426 and S467, have been previously identified as DNA-PK targets in response to DSBs (*22*). Our data demonstrate that, in response to perturbed replication, DNA-PK contributes modestly to phosphorylation at these sites, suggesting that different kinases might target the acidic domain of WRN to modulate its specific functions. Although RPA is important to direct multiple proteins to their genomic DNA substrates in response to replication fork perturbation (*31*), binding of WRN to RPA is dispensable for WRN recruitment to ssDNA in the cell. This finding differentiates the relationship of RPA with WRN from BLM, which requires association with RPA to be localised at ssDNA (*32*). Recently, WRN was shown to cooperate with DNA2 for end-processing of reversed replication forks and during long-range resection that occurs at replication-dependent DSBs induced by CPT treatment (*27*). Interestingly, our data show that RPA-binding is not involved in the WRN/DNA2-dependent end-processing, although RPA-ssDNA complexes are expected to form under these conditions. This would be consistent with RPA-dependent and independent helicase activities of WRN (*33*). However, RPA interacts also with DNA2 and stimulates its function (*34*). Thus, during end-processing at either stalled or collapsed forks, DNA2 would act as an RPA-binding protein for the WRN-DNA2-RPA complex in the same way as described for BLM (*32*).

Similarly, the protective function of WRN against pathological MUS81-dependent DSBs (*25*, *26*) does not require interaction with RPA because the WRN 6A mutant exhibits normal levels of DSBs and RAD51 recruitment after replication fork stalling.

In contrast, our findings provide evidence that RPA-binding by WRN is essential for productive recovery of stalled forks. WRN is copurified with replication factors and defective replication fork progression has been repeatedly reported in absence of WRN in different settings (*25*, *35–39*). Thus, it is tempting to speculate that WRN might perform at least two roles at the perturbed replication fork: end-processing and protection from DSBs in an RPA-independent way or promotion of fork restart in an RPA-dependent manner. Interestingly, association with RPA strongly stimulates WRN helicase activity *in vitro* (*10*, *17*, *19*, *20*). Our observations suggest that WRN helicase activity and WRN binding to RPA act in an epistatic manner. Indeed, either expression of the RPA-defective WRN 6A mutant or pharmacological inhibition of WRN helicase activity in wild-type cells impairs fork restart. WRN, *in vitro*, can also unwind secondary DNA structure, such as hairpins or G4s (*40*, *41*). We observe that impairment of RPA-binding by WRN or inhibition of WRN helicase activity induces G4s accumulation upon fork stalling, suggesting that WRN’s interaction with RPA might render WRN helicase competent for clearance of G4 obstacles and perhaps other secondary DNA structures. Consistent with this idea, loss of WRN sensitizes cells to chromosome breakage at common fragile sites, which are prone to secondary DNA structure formation, and WRN helicase function is important to overcome perturbed replication at these loci (*42–44*). In addition, loss of WRN helicase has been shown to sensitize cells to extended di-nucleotide repeats accumulating in microsatellite unstable cancers, possibly justifying the genetic relationship of synthetic lethality shown by WRN in this background (*8*, *45*, *46*).

It will be interesting to investigate if RPA-binding defective WRN 6A mutant also confers any telomeric phenotype as telomeric DNA is prone to secondary DNA structure formation and WRN is implicated in telomere biology (*47*, *48*). It is worth noting that RPA binding is important also for BLM-mediated fork restart (*32*) suggesting that RPA might be generally required to stimulate RECQ helicases acting at “complex” substrates.

Notably, defective fork restart associated with loss of RPA-binding by WRN also results in the accumulation of regions of ssDNA in the template strand; however, these regions do not arise from repriming by PRIMPOL as shown in the absence of BRCA2 (*49*, *50*). In contrast, our data indicates that these parental ssDNA regions are produced by MRE11 and required for G4 removal *via* the formation of MUS81-dependent DSBs (see model in Figure 8). MRE11 exonuclease activity might be involved in enlarging gaps before MUS81 endonuclease-mediated cleavage of the G4s and, perhaps, to further resect the end of the DSBs introduced at the G4. It is worth noting that although loss of WRN stimulates formation of MUS81-dependent DSBs, this does not occur upon sensitization of common fragile sites (*26*, *51*). As common fragile sites are thought to form secondary DNA structures, this might suggest that the kind of replication perturbation induced at common fragile sites is not resolved by the same mechanism acting at other secondary DNA structures, such as G4s. Interestingly, MUS81 has been involved in the cleavage of G4s at stalled forks and in the removal of secondary DNA structures arising at expanded dinucleotide repeats formed in MMR-deficient cancer cells (*8*, *30*). We observe that MUS81-dependent DSBs are subsequently channelled through a RAD51-dependent post-replication repair as previously shown for some gaps left behind MMS-perturbed forks (*52*, *53*). This pathway is a true salvage mechanism since its abrogation leads to DSBs accumulation (Figure 8).

**Fig. 8.**
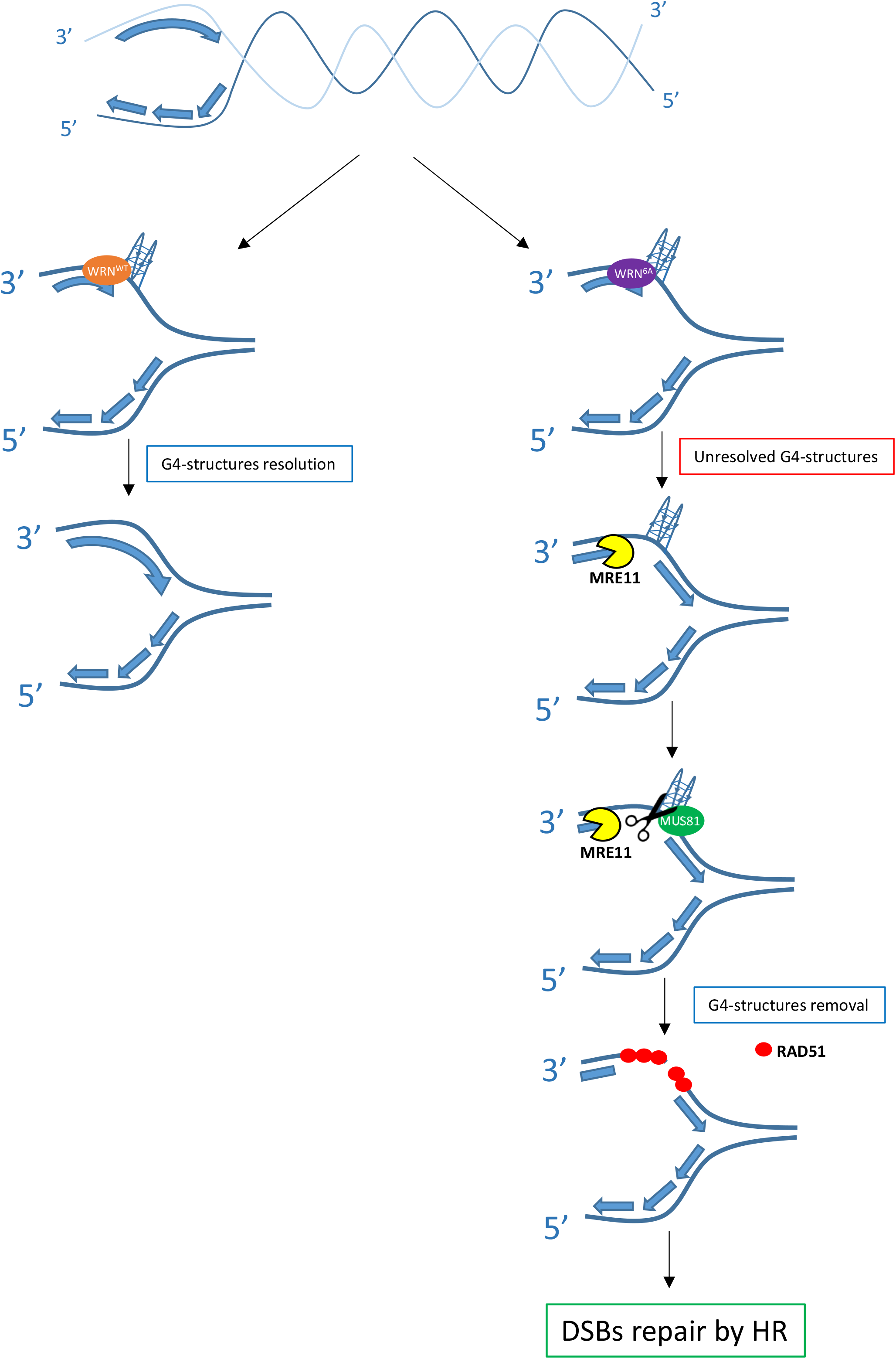
Model of G4s removal when the WRN-RPA interaction is defective. Replication fork stalling occurring near secondary-prone DNA structures, such as guanine-rich regions, can stimulate formation of secondary DNA structures like G4, as depicted. For sake of simplicity, the G4 has been sketched only in the leading strand. These DNA structures require multiple proteins for their resolution, including WRN together with its partner RPA. When WRN cannot properly bind to RPA, these structures persist, and replication is restored downstream leaving a gap in the template. During replication recovery, these gaps are processed by MRE11-exo allowing the MUS81 endonuclease to induce a DSB followed by post-replication repair by RAD51 and “removal” of the secondary DNA structure such as G4s.

In conclusion, we determined that loss of RPA binding to WRN represents a true separation-of-function mutation that interferes with WRN’s cellular functions during the replication stress response or DSBs repair. Because most of the phosphorylation sites are conserved in the mouse Wrn and some also in C. elegans (Supplementary Figure 2), future studies in animal models, in which the RPA-binding ability of WRN is impaired by loss of phosphorylation mutations in the acidic domain, might be useful in elucidating which function(s) of WRN is essential to prevent characteristic phenotypes associated with the accelerated aging of Werner’s syndrome.

## LIMITATIONS OF THE STUDY

Since site-specific mutations abrogating WN-RPA interaction do not exist and deletion of the acidic domain leads to unstable protein, in our study, to address the functional role of the WRN-RPA complex, we used a regulatory mutant of WRN that cannot be phosphorylated by CK2. We show that loss of CK2-dependent phosphorylation leaves unaffected most of the known WRN’s function at perturbed replication forks. However, this mutant is primarily a CK2 unphosphorylable protein, and we cannot rule out that it may be defective in other processes we did not formally test, such as NHEJ or for interactions with other factors outside S-phase. Similarly, although CoIP experiments show the absence of WRN in complex with RPA when CK2-dependent phosphorylation is prevented, a low level of WRN-RPA interaction can be observed at single cell level. Residual interaction with RPA seems to be irrelevant for the fork recovery, however, we cannot rule out that this residual interaction, likely deriving from the secondary binding site in the C-terminal region of WRN, contributes to some WRN function at replication forks.

## MATERIALS AND METHODS

### Cell lines and culture conditions

The SV40-transformed WRN-deficient fibroblast cell line (AG11395) was obtained from Coriell Cell Repositories (Camden, NJ, USA). The AG11395 cell line carries an Arg368 stop mutation in the WRN coding sequence that gives rise to a truncated protein that is degraded and undetectable. AG11395 (WS) were nucleofected with plasmids encoding pCMV-Flag WRN wt and the unphosphorylable (6A) and the phosphomimetic (6D) form of WRN. CK2 phosphorylation mutants were made by replacement of threonine 434, 461 and serine 435, 440, 432 and 467 with alanine or aspartic acid. HEK293T cells were from American Type Culture Collection and they are transfected with the same WRN plasmids.

All the cell lines were maintained in Dulbecco’s modified Eagle’s medium (DMEM) supplemented with 10% FBS with or without tetracycline and incubated at 37 °C in a humidified 5% CO2 atmosphere.

### Chemicals

– Hydroxyurea (HU 98% powder, Sigma-Aldrich) was dissolved in ddH20 and used at 2mM.
– Silmitasertib (CX-4945 Selleck), an inhibitor of CK2 kinase activity, was dissolved in DMSO and a stock solution (500μM) was prepared and stored at –80°C. It was used at 20 μM.
– NU7441 (Selleck), a DNAPKcs inhibitor, was stocked at 1mM in DMSO and used at final concentration of 1 μM.
– Mirin (MRE11i) (Calbiochem), an inhibitor of MRN-ATM pathway, was stocked at 50mM in DMSO and used at 50 μM.
– NSC617145 (Tocris Bio-Techne), an inhibitor of WRN helicase activity, was stocked at 10 mM in DMSO and used at 4 μM.
– HY-128729 (Thermo Fisher), an inhibitor of DNA2 activity, was stocked at 150mM in DMSO and used at 300 μM.
– B02 (553525 Sigma-Aldrich), an inhibitor of RAD51, was dissolved in DMSO a stock solution (37Mm) was prepared and stored at –20°C. It was used at 37 μM.
– 5-iodo-2’-deoxyuridine (IdU) and 5-Chloro-2′-deoxyuridine (CldU) (Sigma-Aldrich) were dissolved in sterile DMEM at 2.5mM and 200mM respectively and stored at –20°C. IdU was used at 100μM for single strand assay and 250μM for fiber assay. CldU was used at 50 μM.
– 5-ethylene-2′-deoxyuridine (EdU) (Sigma-Aldrich) was dissolved in sterile DMSO at 125 mM and stored at –20°C. It was used at 125 μM for 8 mins for SIRF assay.
– Hs MUS81 6 FlexiTube siRNA cat #SI04300877 was stocked at 20 μM and used at 20 nM to knock-down MUS81
– Hs PRIMPOL Silencer select siRNA cat#4427037 was stocked at 20 μM and used at 25 nM to knock-down PRIMPOL

### Nucleofection and Transfection

AG11395 and HEK293T cells were nucleofected and transfected with pCMV Flag WRN wt, pCMV Flag WRN 6A plasmids. For the nucleofection 10μg of DNA were used for 1.8×10^6 cells, with 2 pulses of 950V lasting 2 ms by Invitrogen Neon Transfection system (Invitrogen). After 24hrs in empty medium, cells were placed in 10% FBS medium. 293T cells were transfected with Dreamfect (OzBioscences): 20 μl of Dreamfect was used with 5 μg of DNA, mixed in empty medium for 18 mins. After 24h in empty medium, cells were put in 10% FBS medium.

### Generation of the GST-WRN fragment

DNA sequence corresponding to aa 402-503 (N-WRN) of WRN was amplified by PCR from the pCMV-FlagWRN (wt) plasmid and pCMV-FlagWRN (6D). The PCR product were subsequently purified and sub-cloned into pGEX4T-1 vector (Stratagene) for subsequent expression in bacteria as GST-fusion proteins. The resulting vectors were subjected to sequencing to ensure that no mutations were introduced into the WRN sequence in the plasmid used for transforming BL21 cells (Stratagene). Expression of GST and GST-fusion proteins were induced upon addition of 1 mM isopropyl-D-thiogalactopyranoside (IPTG) for 2 hrs at 37°C. GST, and GST-N-77 WRN were affinity-purified using glutathione (GSH)-magnetic beads (Promega). Fragment purification levels were assessed by SDS-PAGE followed by Coomassie staining.

### Purification of recombinant FLAG-WRN

High titer virus expressing Flag-WRN was used to infect Hi5 insect cells (Thermo Fisher Scientific) at an MOI of approximately 10. Cells were harvested 48 hours later and placed in –80° C until lysed. Cell pellet containing approximately 1.2 x 108 cells was resuspended in 10 ml of Lysis Buffer (50 mM Tris pH 7.4, 150 mM NaCl, 0.4% NP40, 10% glycerol, 5 mM BME, and Complete Ultra Protease Inhibitors (Roche)), vortexed and rotated at 4° C for 45 mins. The lysates were centrifuged at 20,000 RPM for 10 minutes and the supernatant was passed through a 0.45 µm PVDF filter. Each clarified lysate was passed twice through a Ni2+-charged 1 ml HiTrap Chelating HP column (GE Healthcare Life Sciences) which had been equilibrated in Buffer TN (50 mM Tris pH 7.4, 150 mM NaCl, 10% glycerol, 5 mM BME, protease inhibitors) with 10 mM imidazole. 5 ml washes with TN buffer containing 10 mM, 20 mM, and 40 mM imidazole each were performed followed by elution with TN buffer containing 400 mM imidazole. The eluted protein was pooled and incubated with TEV protease for 16 hrs at 4° C to cleave the 6×His tag off the protein. The protein was dialyzed into NETN-500 Buffer (50 mM Tris pH 7.4, 500 mM NaCl, 0.5% NP40, 1 mM EDTA) using a Amicon Ultra 100 kD cutoff centrifugal filter (EMD Millipore). The retained sample was applied to 250 ul of packed M2 anti-FLAG beads (Sigma) which had been equilibrated in NETN-500 buffer. The beads were washed twice with 5 ml of NETN-500 buffer and the WRN protein was eluted with 3X FLAG peptide twice in 500 µl Storage Buffer (100 mM Tris pH 8.0, 400 mM NaCl, 10% glycerol, 5 mM BME). Eluted protein was concentrated and dialyzed against storage buffer in the absence of FLAG peptide and frozen at –80° C.

### Helicase assays

The 25 base pair forked duplex DNA substrate with 35 nt tails (X12-1-rCCTRL: 5’-TTTTTTTTTTGACGCTGCCGAATTCTGGCTTGCTAGTACGCGAGCTCCATCGTTGACCCT-3’, X12-2-rCCTRL: 5’-AGGGTCAACGATGGAGCTCGCGTACGTTTGGTGTAATCGTCTATGACGTTTTTTTTTTT –3’) was prepared as previously described (*54*). Briefly, 0.5 µg of purified recombinant WRN protein was treated with Lambda Protein Phosphatase (LPP, New England Biolabs) in 1X PMP buffer (50 mM HEPES pH 7.5, 100 mM NaCl, 2 mM DTT, 0.01% Brij 35, 10 µl reaction) for 30 minutes at 30°C. Halt Phosphatase Inhibitor Cocktail (Thermo Scientific) was added to 1X final concentration, followed by Casein Kinase II (New England Biolabs) in 1X PK buffer (50 mM Tris-HCl pH 7.5, 10 mM MgCl2, 0.1 mM EDTA, 2 mM DTT, 0.01% BriJ 35, 30 µl reaction) for 30 minutes at 30°C. Untreated, CK2-and/or LPP-treated WRN protein (concentrations indicated in figure legend), or storage buffer were incubated with 0.5 nM forked DNA substrate in 20 µl reactions containing 30 mM HEPES pH 7.4, 40 mM KCl, 100 µg/ml BSA, 8 mM MgCl2, 5% glycerol, and 2 mM ATP for 15 minutes at 37° C in the presence or not of RPA heterotrimer. Reactions were stopped by adding 20 µl of 9 mM EDTA, 0.6% SDS, 0.04% bromphenol blue, 0.04% xylene cyanol, and 25% glycerol containing a 10-fold excess of the labeled oligo without the radiolabel. The heat-denatured sample was boiled for 5 minutes at 95° C.

### In vitro Kinase assay

For kinase assay, 1 μg of immunopurified GST-tagged WRN fragment was phosphorylated in vitro by Casein Kinase II (New England Biolabs) in the presence of 1X NEBuffer (50mM Tris-HCl, 10mM MgCl2, 0.1 mM EDTA, 2mM DTT, 0.01% BriJ 35) and 200μM ATP for 30 minutes at 30°C.

### Pulldown Assay

GST and GST-WRN fragments (phosphorylated or not) were incubated with 300ng of 293T cell extracts. After 16hrs of incubation, fragments were separated from the beads and RPA32 interaction with WRN fragments were measured with densitometric analysis by WB using rabbit anti-GST (Calbiochem), rabbit anti-p440WRN (Abgent) and mouse anti-RPA34-20 (Millipore).

### Immunoprecipitation and Western blot analysis

Immunoprecipitation experiments were performed using 3×106 293T cells. IP buffer (0.5% Triton X-100, 50mM Tris HCl pH 8.0, 150 mM NaCl, EGTA 1 mM) supplemented with phosphatase, protease inhibitors and benzonase was used for cells lysis. Two mL of lysate were incubated overnight at 4°C with 20 μl of Anti-Flag M2 magnetic beads (Sigma) or Anti-RPA32 conjugated Dynabeads (2μg of MABE285 anti-RPA34-20 mouse (Millipore) with 40μl of Dynabeads protein G (Invitrogen). After extensive washing in IP buffer, proteins were released in 2X Laemmli buffer buffer and subjected to Western blotting.

Blots were incubated with primary antibodies: rabbit anti-FLAG (Sigma-Aldrich); rabbit anti-RPA70 (Genetex); mouse anti-RPA34-20 (Millipore); rabbit anti-pS440/467WRN (Abgent; custom-made); rabbit anti-Lamin B1 (Abcam); rabbit anti-RAD51 (Santa Cruz); rabbit anti-GST (Calbiochem). Blots were detected using the Western blotting detection kit WesternBright ECL (Advansta) according to the manufacturer’s instructions. Quantification was performed on scanned images of blots using Image Lab software, and values shown on the graphs represent normalization of the protein content evaluated through Lamin B1 or Immunoprecipitated protein immunoblotting.

### Mass Spectrometry Analysis

Identification of phosphopeptides was performed as already described (*55*). Briefly, purified proteins were in gel-digested with trypsin, phosphopeptides enriched by IMAC following the manufacturer’s guidelines (Phosphopeptide Enrichment Kit; Pierce) and mass spectrometry analysis performed with a MALDI-TOF Voyager DE-STR (Applied Biosystems, Foster City, CA, USA) in positive reflectron mode, using phospho-DHB as matrix. MS spectra were processed with DATA EXPLORER (Applied Biosystems) and GPMAW software for peak to sequence assignments. To confirm the attribution of relative peaks to mono-, di-and tri-phosphorylated peptides, alkaline phosphatase treatment was performed on-probe as already described (*56*).

### Chromatin isolation

To isolate chromatin, cells were resuspended in buffer A (1M Hepes pH 7.9, 1M HCl, 100mM MgCl2, glycerol, sucrose, sodium fluoride, 1 mM DTT, protease inhibitors, phosphatase inhibitors). Triton X-100 (0,1%) was added, and the cells were incubated for 5 min on ice. Nuclei were collected by low-speed centrifugation (4 min, 4000 rpm, 4°C). Nuclei were washed once in buffer A, and then lysed 10 min in buffer B (3 mM EDTA, 0,2 mM EGTA, 1 mM DTT, protease inhibitors, phosphatase inhibitors). Insoluble chromatin was collected by centrifugation (4 min, 4500 rpm, 4°C), washed once in buffer B, and centrifuged again under the same conditions. The final chromatin pellet was resuspended in 2X Laemmli buffer and sonicated for 15 sec in a Tekmar CV26 sonicator using a microtip at 25% amplitude. Lastly, the lysates were subjected to Western blot analysis. As a specific loading we probed blots for Lamin B1, or H3 histone which are proteins exclusively found in the chromatin fraction.

### Single-stranded DNA detection and immunofluorescence assay

Cells were cultured onto 22×22 coverslip in 35mm dishes. To detect nascent single-stranded DNA (ssDNA), after 24 hrs, the cells were labelled for 15 min before the treatment with 100μM IdU (Sigma-Aldrich), cells were then treated with HU 2mM for different time points. Meanwhile, to detect parental ssDNA, the cells were labelled for 24 hrs before 2 hrs of fresh medium. After the release, cells were treated with HU. Next, cells were washed with PBS, permeabilized with 0.5% Triton X-100 for 10 min at 4°C and fixed with 2% sucrose, 3%PFA. For ssDNA detection, cells were incubated with primary mouse anti-IdU antibody (Becton Dickinson) for 1h at 37°C in 0.1% saponine/BSA in PBS, followed by Alexa Fluor488-conjugated goat-anti-Mouse, and counterstained with 0.5μg/ml DAPI. Instead, to detect RPA32 and WRN, cells were incubated with specific primary antibody: rabbit anti-WRN (Abcam) and mouse RPA34-20 (Millipore) for 1h at 37°C in 0.1% saponine/BSA in PBS followed by Alexa Fluor 594 Anti-Rabbit or Alexa Fluor 488 Anti-Mouse. For immunofluorescence of G-quadruplex structures (G4s), cells grown on glass coverslips were fixed with ice-cold 80% methanol in PBS for 15 min at –20°C, then washed two times in PBS. Next, cells were blocked with 10% FBS/PBS for 1 h and incubated with the anti-G-quadruplex antibody (BG4, Sigma-Aldrich, 1:200) overnight at 4°C. Slides were analyzed (at 40×) with Eclipse 80i Nikon Fluorescence Microscope, equipped with a Virtual Confocal (ViCo) system. Fluorescence intensity for each sample was then analyzed using ImageJ software.

### In situ proximity ligation assay (PLA)

Cells were cultured onto 8-well Nunc chamber-slides. The *in situ* proximity ligation assay (PLA) in combination with immunofluorescence microscopy was performed using the Duolink Detection (Merck) or the NaveniFlex (Navinci diagnostics) Kit with anti-Mouse PLUS and anti-Rabbit MINUS PLA Probes, according to the manufacturer’s instructions. To detect proteins, we used rabbit anti-WRN (Abcam), rabbit anti-Flag (Sigma-Aldrich), mouse anti-RPA34-20 (Millipore), rabbit anti-RAD51 (Santa Cruz) and mouse anti-IdU antibody (Becton Dickinson) antibodies.

### DNA fiber analysis

Cells were pulse-labelled with 50 μM CldU and then labeled with 250 μM IdU at the times specified, with HU treatment as reported in the experimental schemes: to study DNA degradation cells were labelled for 20 min of CldU, then after 2 washes in PBS, they were labelled for 20 min of IdU. After another 2 washes, cells were treated with 2mM HU at different times. Meanwhile, to study the ability to recover from replicative stress, cells were labelled for 20 min before the HU treatment and during the treatment itself. After 2 washes, cells were labelled for 20 min with IdU. DNA fibers were prepared, spread out and immunodecorated following primary antibodies with rat anti-CldU/BrdU (Abcam) and mouse anti-IdU/BrdU (Becton Dickinson). Images were acquired randomly from fields with untangled fibres using Eclipse 80i Nikon Fluorescence Microscope, equipped with a Virtual Confocal (ViCo) system. The length of labeled tracks were measured using the ImageJ software. A minimum of 100 individual fibres were analyzed for each experiment.

### Neutral Comet assay

After treatment, cells were embedded in low-melting agarose and spread onto glass slides. After an electrophoretic run of 20’ (6-7 A, 20V), cells were fixed with methanol. DNA was stained with 0.1% GelRed (Biotium) and examined at 20× magnification with an Olympus fluorescence microscope. Slides were analysed with a computerized image analysis system (CometScore, Tritek Corp.). To assess the amount of DNA DSB breaks, computer generated tail moment values (tail length × fraction of total DNA in the tail) were used. Apoptotic cells (smaller comet head and extremely larger comet tail) were excluded from the analysis to avoid artificial enhancement of the tail moment. A minimum of 150 cells were analyzed for each experiment.

### Statistical analysis

All the statistical analysis was performed using GraphPad Prism 9 software. Frequency distributions of DNA track length and ratios were determined with GraphPad Prism 9 software after the quantification of the tract lengths using ImageJ. Mann-Whitney and Kruskal-Wallis tests coupled with ad-hoc Dunn test for false-discovery rates were used for statistical analyses when comparing two and more than two variables, respectively. In all graphs, P < 0.05 was considered significant for frequency distribution. When data are not presented as scatter plots they are shown as the mean of independent experiments.

## ACKNOWLEDGMENTS

We want to thank Prof. Achille Pellicioli for helpful discussion and advise. We thank Dr. Giuseppe Leuzzi for the help with initial characterisation of fork progression phenotype in HEK293T cells expressing the WRN mutant. This work was supported by investigator grants from Associazione Italiana per la Ricerca sul Cancro (AIRC) to P.P. (IG n. 21428) and to A.F. (IG n. 19971), and in part by NIH (grant n. 1R01DE029471-01A1) and the Intramural Research Program, National Institute on Aging, NIH.

## AUTHORS CONTRIBUTION

A.N. performed the characterisation of WRN phosphorylation mutant in vitro and in vivo, and the majority of the functional analysis of the CK2-phosphorylation WRN mutants. V.P. performed the analysis of end-resection at CPT-induced DSBs and generated the acidic-domain deleted WRN mutant. F.D.F contributed to the analysis of DNA damage in cells expressing the CK2-unphosphorylable WRN mutant. P.P. performed radioactive kinase assays and kinase assays for MS/MS. F.F. performed MS/MS analysis and phosphorylation sites identification. M.S. performed the purification of WRN fragments. P.V. performed the analysis of G4 accumulation. J.A.S. and T.K. performed WRN purification and enzymatic assays. M.C. supervised the MS/MS analysis. V.P., M.S., and S.R. analysed data and contributed to designing the experiments. All authors contributed to design experiments and analyze data. R.M.B., P.P. and A.F. supervised the project and wrote the paper. All authors contributed to revise the paper.

