## Supplementary Figures and Legends for "Phosphorylation-Dependent Association of WRN with RPA is Required for Recovery of Replication Forks Stalled at Secondary DNA Structures"

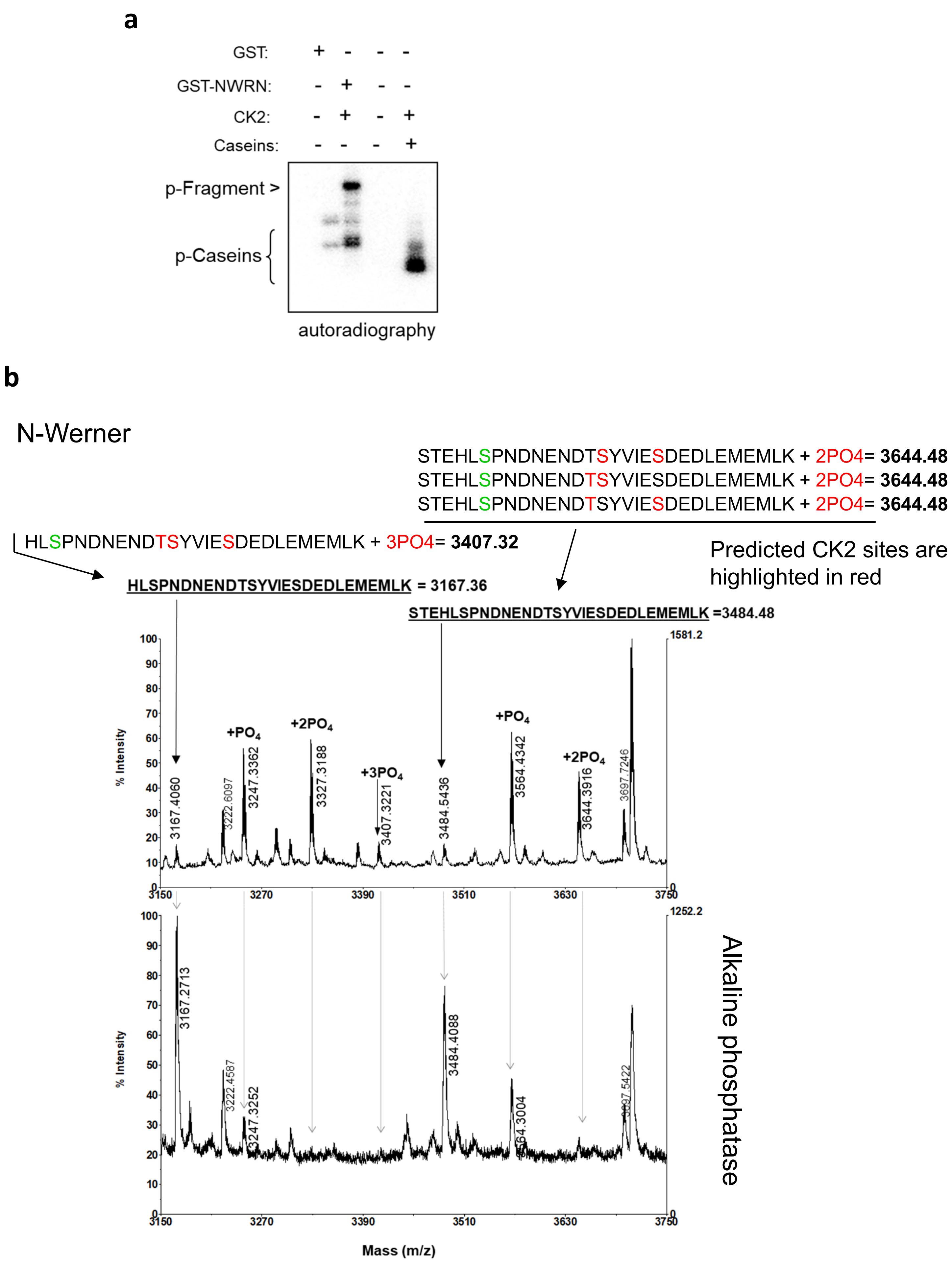


Supplementary Fig. 1

**a**

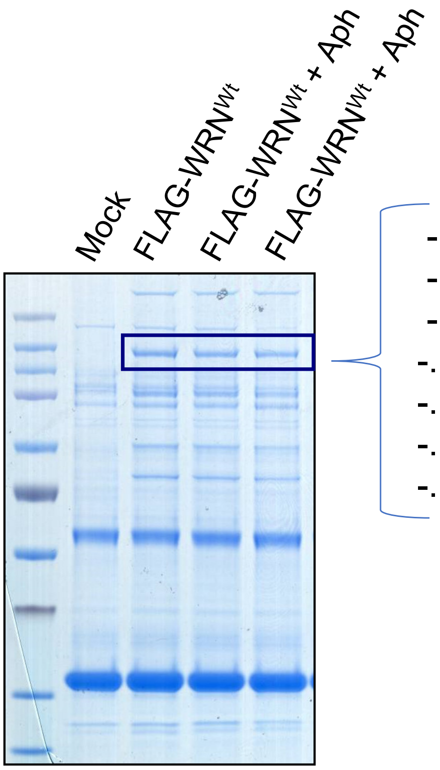


**b**

-.HLSPNDNENDTSYVIE**S#**DEDLEMEMLK.-
-.HLSPNDNENDT**S#**YVIESDEDLEMEMLK.-
-.HLSPNDNEND**T#**SYVIESDEDLEMEMLK.-
-.HLSPNDNEND**T#**SYVIE**S#**DEDLEMEMLK.-
-.HLSPNDNENDT**S#**YVIE**S#**DEDLEMEMLK.-
-.HLSPNDNENDTS**Y#**VIE**S#**DEDLEMEMLK.-
-.HLSPNDNEND**T#**S**Y#**VIESDEDLEMEMLK.-

**c**

```
gi|110735439|ref|HUMAN        TEHELQILEQQSQEEYLSDIAYKSTEHLSPNDNENDTSYVIESDEDLEME 447
gi|170763504|ref|MOUSE        SENELQDLEQQAKEEKYNDVSHQLSEHLSPNDDENDSSYIIESDE----- 435
gi|118089996|ref|Chicken      NEQELQILERQAAEEVLN---KAVPEEECPTDNDQNTSCIIESDE----- 414
gi|29428101|sp|Xenopus        SEEELYMMEREDDKKQTN-PDYKLNKDSCDTNEEKDMSYVIESDE----- 409
                              .*.**  :*::  ::  .       :. . .::::: * :*****

gi|110735439|ref|HUMAN        MLKHLSPNDNENDTSYVIESDEDLEMEMLKSLENLNSGTVEPTHSKCLKM 497
gi|170763504|ref|MOUSE        ----------------------DLEMEMLKSLENLNSDVVEPTHSTWLEM 463
gi|118089996|ref|Chicken      ----------------------ELEMEMLKSLEDVDNSKEVPTERESSKA 442
gi|29428101|sp|Xenopus        ----------------------DFDSEIIKSLEDLDNSTEEALGTGVPQA 437
                                                    ::: *::****:::..   .      :
```

Supplementary Fig. 2

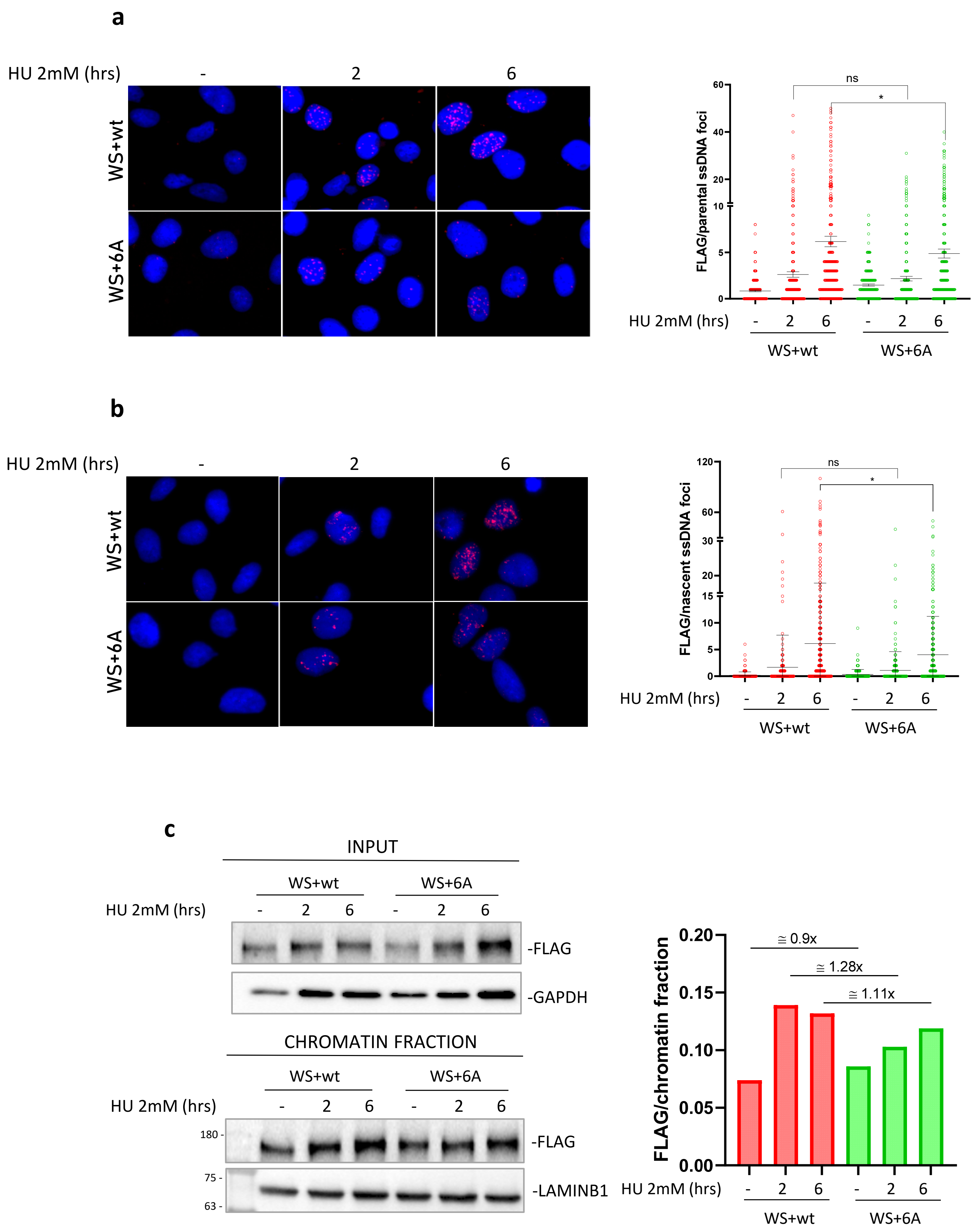


Supplementary Fig. 3

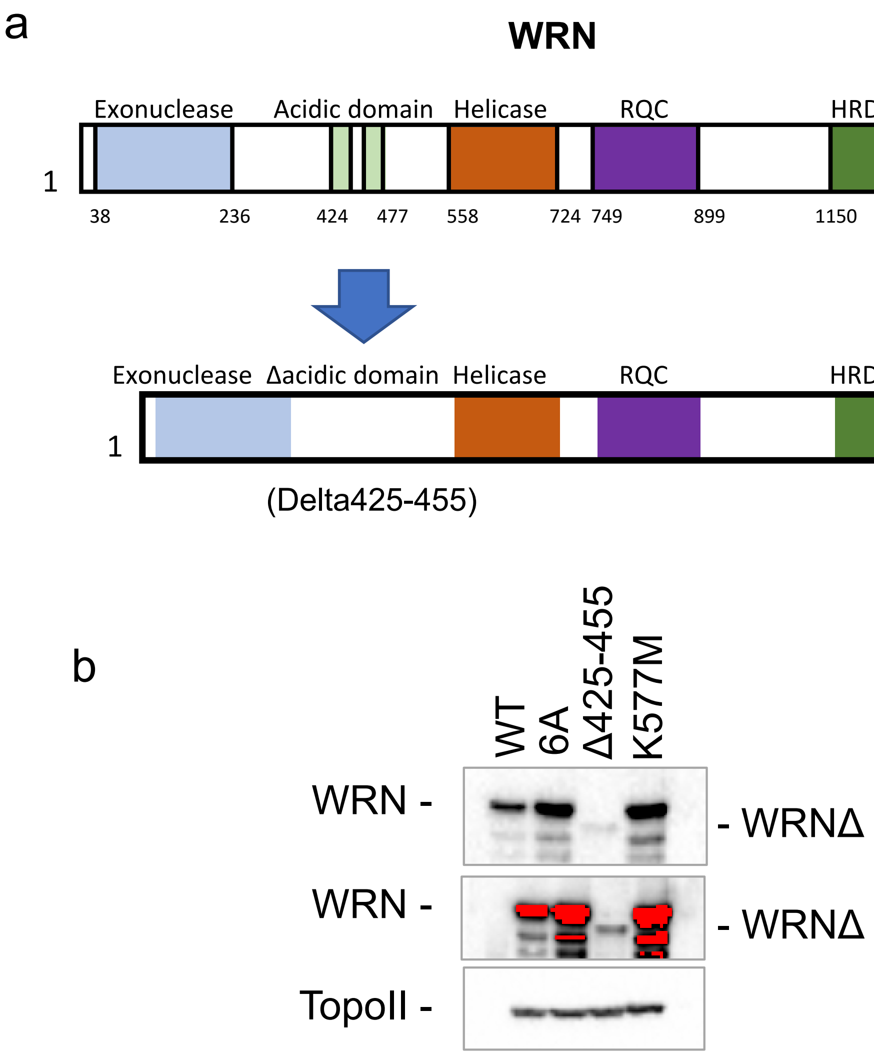


Supplementary Fig. 4

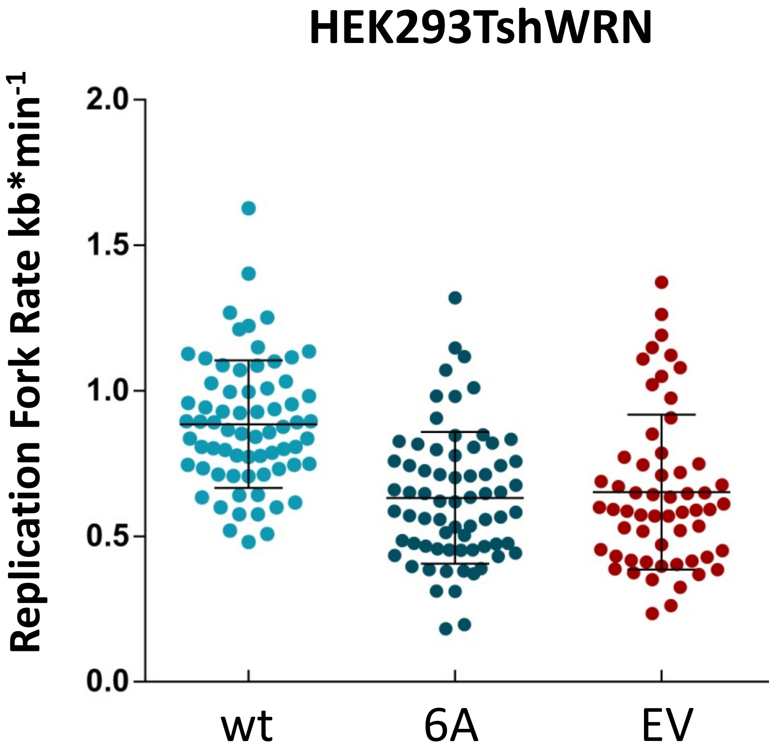


Supplementary  Fig. 5

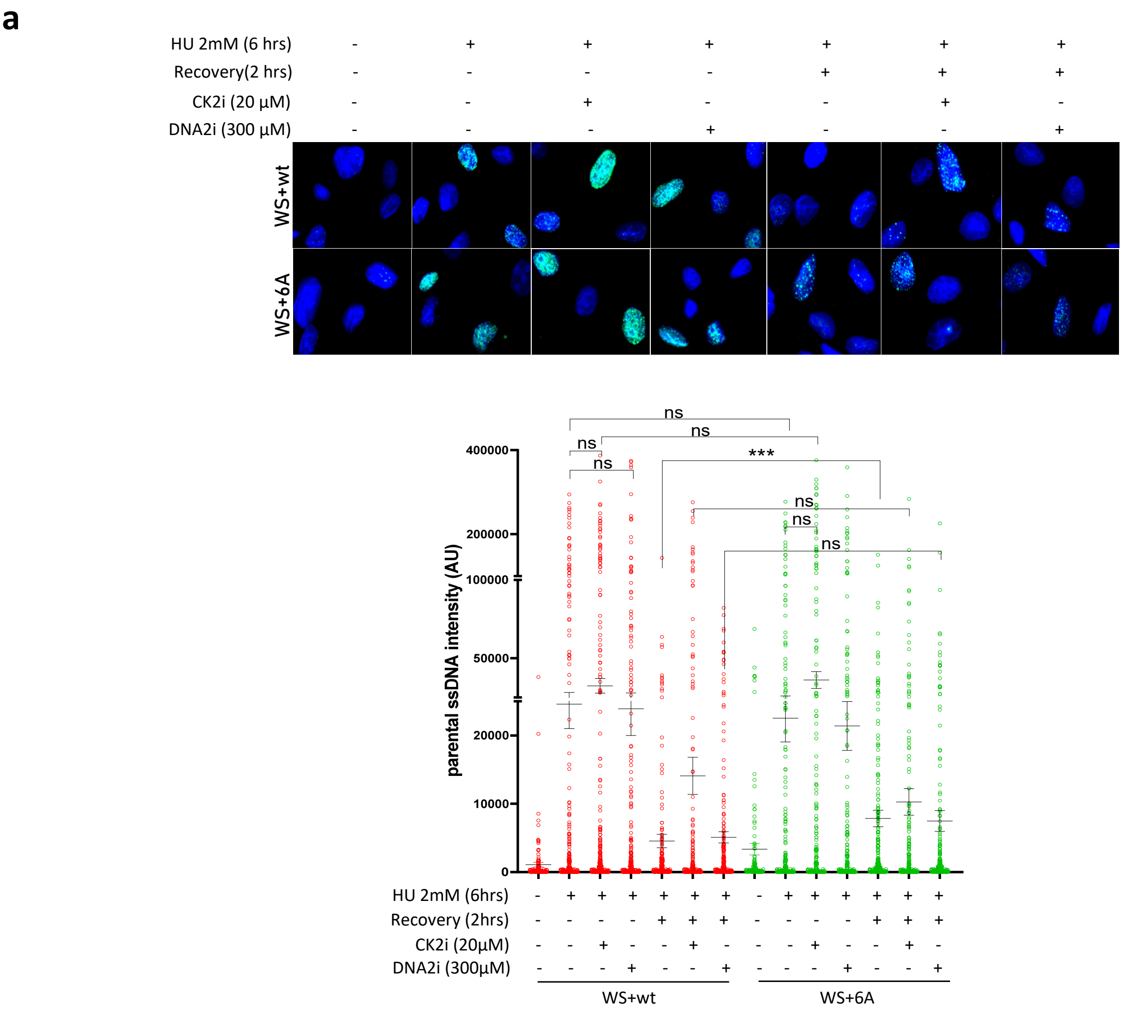


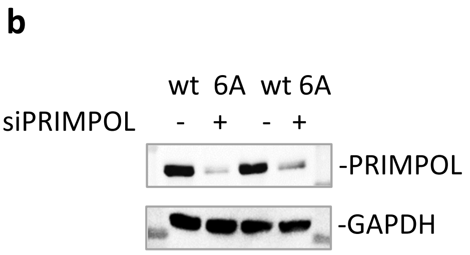


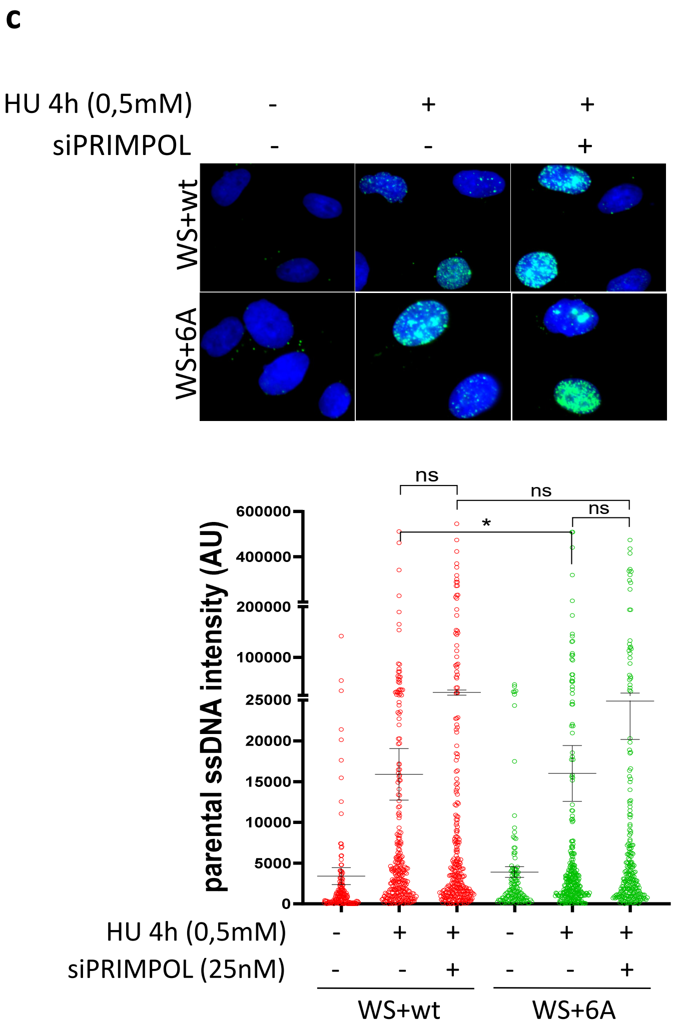


Supplementary Fig. 6

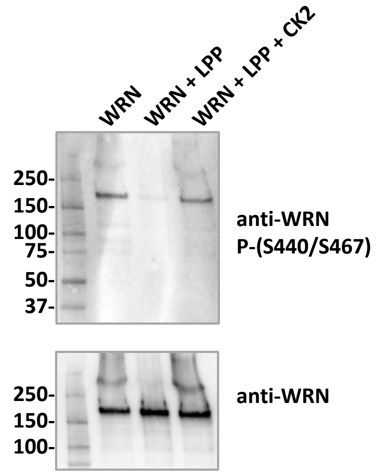


**Supplementary Fig. 7**

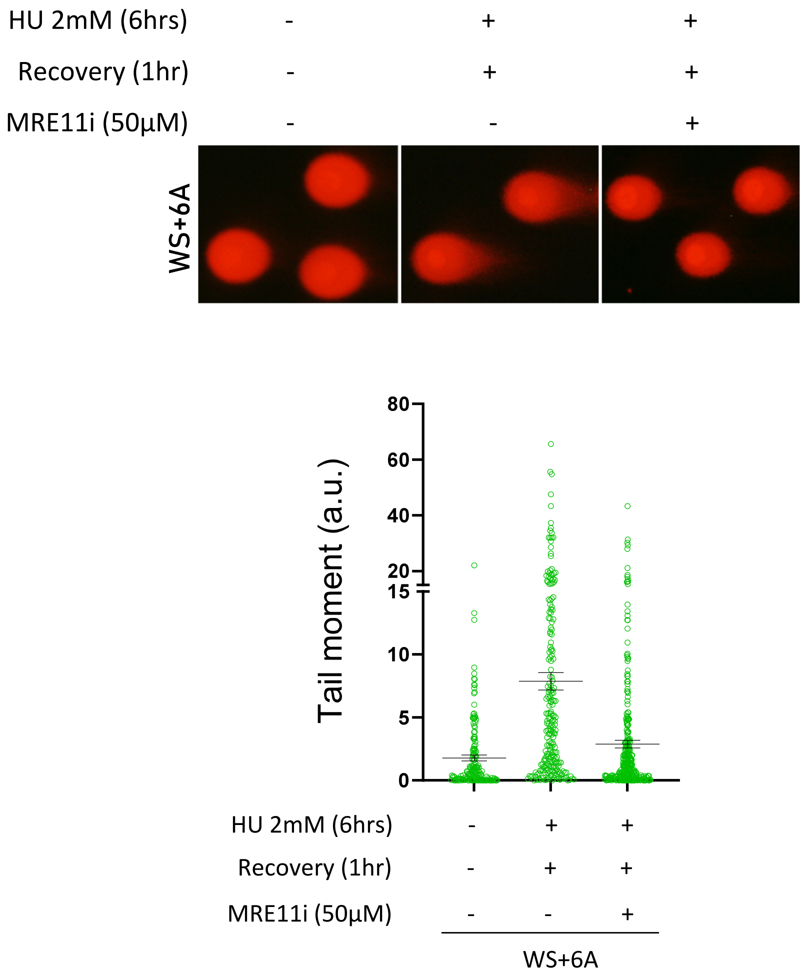


**Supplementary Fig. 8**

SUPPLEMENTARY FIGURES LEGENDS

**Fig. S1. CK2 targets multiple sites in an N-terminal WRN fragment containing the acidic domain.** A) The 1-505 N-terminal region of WRN was purified from bacteria as GST-fusion protein and used as a substrate for in vitro kinase assay. Autoradiography shows phosphorylation of the fragment by CK2. A duplicate, non-radioactive gel, was used in B) for the analysis by mass spectrometry.

**Fig. S2. The acidic domain of WRN is phosphorylated by CK2 in the cell in response to replication arrest.** A) Flag-WRN was immunoprecipitated from HEK293T cells transfected with FLAG-WRNwt and treated with 1 µM aphidicolin for 8h as indicated. The bands corresponding to untreated and the two treated duplicate samples were excised and phosphorylation analysed by mass spectrometry. The identified phosphorylated sites are denoted with # and listed in B). C). Alignment of sequences corresponding to the first repeat of the duplicated region of WRN containing the CK2 sites with sequences from other WRN orthologues from different vertebrates.

**Fig.S3. $WRN^{6A}$ recruitment at stalled or reversed forks is not impaired.** A) Analysis of WRN recruitment at parental ssDNA by *in situ* PLA using FLAG and IdU antibodies. WS cells nucleofected with Flag-$WRN^{wt}$ or Flag-$WRN^{6A}$ were treated as indicated. The graph shows individual values of PLA spots. Representative images are shown. Bars represent mean ± S.E. B) Analysis of WRN recruitment at nascent ssDNA by *in situ* PLA using FLAG and IdU antibodies. The graphs show individual values of PLA spots. Representative images are shown. Bars represent mean ± S.E. Statistical analyses were performed by Kruskal-Wallis test (ns = not significant; *$P<0.05$; Where not indicated, values are not significant). C) Analysis of WRN recruitment in chromatin by cell fractionation. WS cells expressing Flag-tagged $WRN^{wt}$ or $WRN^{6A}$ were treated as indicated and chromatin fraction analysed by Western Blot. The graph shows the quantification of the normalised amount of WRN(Flag) in chromatin from the representative blot.

**Fig.S4. Deletion of large part of the acidic domain of WRN leads to unstable protein.** A) Schematic representation of WRN protein and its truncated version without acidic domain (ΔWRN-Δ425-455). B) Western Blotting analysis of WRN expression in WS cells nucleofected with $WRN^{wt}$, $WRN^{6A}$, ΔWRN and K577M (WRN helicase inactive mutant). Truncated version of WRN is expressed at extremely low levels when compared to wild-type and other mutant forms of WRN.

**Fig.S5. WRN$^{6A}$ nucleofected cells show less fork restart and slower start recovery.** A) Analysis of fork speed for untreated HEK293T cells expressing the WRN6A mutant (6A), WRNwt (wt) or empty-vector (EV). The graph shows individual values of fork rate as extrapolated from the measurement of the IdU tract length in dual CldU/IdU labelled fibers.

**Fig.S6. CK2-dependent WRN phosphorylation is not related with DNA2-dependent end-resection or DNA repriming.** A) Analysis of parental ssDNA in WRN$^{wt}$ and WRN$^{6A}$ nucleofected cells treated with 2 mM HU for 6 hours and recovered with or w/o CK2 or DNA2 inhibitor. The graph shows individual values of total nuclear IdU foci intensity. Bars represent mean ± S.E. Statistical analyses were performed by ANOVA (ns = not significant; *P<0.05; ***P< 0.001). B) Analysis of PRIMPOL downregulation by WB. C) Analysis of parental ssDNA in WRN$^{wt}$ and WRN$^{6A}$ nucleofected cells treated with 0.5 mM for 4 hours with or w/o prior depletion of PRIMPOL. The graph shows individual values of total nuclear IdU foci intensity. Bars represent mean ± S.E. Statistical analyses were performed by Kruskal-Wallis (ns = not significant; *P<0.05; Where not indicated, values are not significant). Representative images are shown in the panels.

**Fig. S7. Recombinant WRN purified from insect cells is phosphorylated in the acidic domain**. Western blot of untreated, CK2- and/or Lambda protein phosphatase (LPP)-treated WRN purified from insect cells. Each preparation corresponds to the batch used in helicase assays shown in Fig. 5.

**Fig.S8. MRE11 activity is required for DSBs accumulation in WRN6A mutant.** Analysis of DSBs by neutral Comet assay. WS cells nucleofected with Flag-WRN6A were treated with HU and recovered for 1 hour in the presence or absence of the MRE11i Mirin. The graph shows individual tail moment values (n=2). Bars represent mean ± S.E. Representative images are shown. Statistical analyses were performed by Student's t-test (**P< 0.01. ****P<0.0001).
